# Single cell-resolution *in situ* sequencing elucidates spatial dynamics of multiple sclerosis lesion and disease evolution

**DOI:** 10.1101/2023.06.29.547074

**Authors:** C. M. Langseth, P. Kukanja, L. A. Rubio Rodríguez-Kirby, E. Agirre, A. Raman, C. Yokota, C. Avenel, K. Tiklová, A. O. Guerreiro-Cacais, T. Olsson, M. M. Hilscher, M. Nilsson, G. Castelo-Branco

## Abstract

Multiple sclerosis (MS) is a neurological disease characterised by multifocal lesions and smouldering pathology. While single-cell analyses have provided insights into neuropathology, cellular processes underlying MS remain poorly understood. We modelled the cellular dynamics of MS by examining temporal and regional rates of disease progression in the experimental autoimmune encephalomyelitis (EAE) mouse model. By performing single-cell spatial expression profiling using *In situ* sequencing, we annotated disease neighbourhoods during lesion evolution and found centrifugal propagation of active lesions. We demonstrated that disease-associated (DA) glia are dynamic and induced independently of lesions, preceding their formation. Single-cell spatial analysis of human archival MS spinal cord confirmed differential distribution of DA-glia, enabled deconvolution of active and inactive lesions into sub-compartments, and identification of new lesion areas. By establishing a spatial resource of mouse and human MS neuropathology at a single-cell resolution, our study unveils the intricate cellular dynamics underlying MS disease evolution.

## INTRODUCTION

Multiple sclerosis (MS) is a complex inflammatory and neurodegenerative demyelinating disorder affecting around 2.8 million individuals worldwide. Its primary symptoms include impaired mobility and cognition, fatigue, numbness, vision problems, depression, and poor coordination. The cause of MS is unclear and multifactorial, involving interaction between genetic predisposition and lifestyle/environmental factors such as Epstein-Barr virus infection, smoking, and obesity, among others^1^. Of these, Epstein-Barr virus infection may constitute a prerequisite to developing MS^2^. Most patients initially develop a relapsing-remitting course of the disease in their late 20s, transitioning later to a secondary progressive stage. Other patients develop directly a primary progressive course of the disease, with a continual and worsening accumulation of the symptoms without remissions^3^.

MS affects various anatomical regions of the central nervous system (CNS), including white matter (WM) and grey matter (GM) of the brain and the spinal cord^4^. Multifocal tissue compartments, defined by inflammation, gliosis, demyelination, and/or neurodegeneration, are recognised as lesions, with their location often, but not always, reflecting clinical symptoms^5, 6^. Nevertheless, subsets of patients have lesions predominantly or limited to certain anatomical regions, such as the spinal cord^7^, suggesting that the tissue microenvironment plays a role in disease specification. Not only lesion distribution but also lesion composition highly varies between patients and within affected individuals. Neuropathological analysis delineating pathological processes in *post mortem* MS tissue has led to several proposed strategies for staging lesions into different categories^8^, with the most updated approach stratifying lesions into active, mixed active/inactive, and inactive with or without ongoing demyelination, or remyelination^9^. In demyelinating lesions, the lesion area with its corresponding status of (de)myelination and activity is, in principle, determined upon staining against myelin, in combination with immunohistochemical evaluation of microglia and macrophages that dynamically change over lesion development. Lesion classification strategies thus seem to gravitate towards simple but broadly reproducible histological methods that can be achievable in most clinical and preclinical research laboratories, and lesion types easily correlated with clinical evaluation or imaging. In a preclinical setting, such strategies provide a broad overview of lesion-type heterogeneity. In turn, a better understanding of cellular and molecular lesion composition would aid in refining current lesion stratification approaches, discovering new biomarkers, and developing more targeted treatments.

Currently, focal inflammation, driven by immune infiltration of the parenchyma, meninges and perivascular spaces, is thought to govern the neuropathology in MS^4^. It has also been suggested that the location and composition of the infiltrates lead to different types of inflammation, which might be associated with different disease courses^10^. For example, parenchymal infiltration would give rise to active lesions, while accumulation of B- and T-cells in meninges and perivascular spaces would give rise to subpial lesions and widespread diffuse neurodegeneration^10^. However, the current human MS neuropathological characterisation relies on *post mortem* tissue collected at the end stages of the disease. Therefore, such analysis might only reflect the final outcomes of a series of disease-related events that occurred during disease onset and progression rather than processes that trigger and contribute to MS progression. A chronological characterisation of lesions and different inflammatory and overall neuropathological events would be essential to elucidate how the disease develops in different cohorts of MS patients.

The rise of single-cell and single-nuclei RNA sequencing (RNA-seq) technologies has uncovered differential transcriptional responses within populations of neurons^11^, glia^12–16^, and vascular cells^17, 18^ in human archival MS tissue and its most commonly used mouse model, experimental autoimmune encephalomyelitis (EAE)^19^. Surprisingly, not only microglia but also astrocytes and oligodendroglia have emerged as highly transcriptionally heterogeneous cellular populations with distinct disease-associated (DA) states arising in response to inflammation^11–15, 20–22^. In particular, DA-oligodendroglia were first described in mouse MS models, with oligodendrocyte precursor cells (OPCs) expressing major histocompatibility complex (MHC) class I and II^14, 23^. DA-OPCs were shown to phagocytose myelin and to be able to activate CD4 and CD8 T-cells by antigen presentation^14, 23, 24^. Subsequent studies demonstrated the prevalence of DA-OPCs and DA mature oligodendrocytes (MOL) also in ageing and other neurodegenerative pathologies, including Alzheimer’s disease^20, 25–29^. However, the functions of DA-glia in the context of MS and other diseases are poorly understood due to the lack of knowledge on their *bona fide* abundances, spatial distributions, and cell-to-cell interactions throughout the disease. Moreover, whether the newly uncovered disease-associated (DA) cellular subtypes play detrimental or neuroprotective roles in MS aetiology and progression remains unclear. Determining where in the CNS these cellular subtypes arise and localise at different stages of the disease is key to understanding their functions.

To bridge this gap, spatial transcriptomics (ST) emerged and was recently implemented in *post mortem* MS brain tissue samples^30, 31^. Although ST is an unparalleled platform to unravel tissue architecture and molecular pathways altered in MS^30, 31^, the current resolution (50μm) does not directly provide RNA counts for single cells. Inferring individual cells in ST relies on integration with scRNAseq datasets for cellular deconvolution^32^, which does not directly translate into the native positions of cells, making the investigation of cellular composition and proximity-based interactions challenging.

Here we use *In situ* Sequencing (ISS)^33–35^ to characterise at a single-cell level the spatio-temporal dynamics of major cellular subtypes in EAE and MS spinal cord archival tissue. We utilise EAE to model the early, acute and resolving inflammation-driven pathology of MS and correlate our findings with MS *post mortem* cervical spinal cord sections. Given that the disease aetiology and progression in EAE occur in a well-defined temporal and spatial fashion, this approach allowed us to map for the first time the dynamics of cellular interactions that potentially also mediate these processes in human MS, establishing the sequence of cellular events underlying lesion development and progression with unprecedented resolution. Analysis of cellular neighbourhoods revealed the formation of five distinct inflammatory tissue compartments during the EAE disease course, which carried regional preferences and profound compositional differences. At the peak of the disease, brain immune-infiltrated compartments arose in the medial corpus callosum and ventral regions of the brain and were accompanied by localised transition of glia into disease-associated (DA) states. In contrast, inflammatory and lesion load was pronounced within the spinal cords and increased from cervical to lumbar, with ventrolateral regions exhibiting more complex, compartmentalised lesions. Additionally, spatio-temporal dynamics of the annotated compartments led to a model of centrifugal lesion propagation in EAE. Surprisingly, at peak EAE, the DA-glia were abundantly distributed throughout the spinal cord parenchyma and diminished with proximity to the lesion cores. In contrast, spinal cords from the late stages of EAE were predominantly populated with homeostatic macroglia, while DA-oligodendroglia and DA-astrocytes were limited to the WM and were further enriched in lesions. A large fraction of microglia retained the DA signatures throughout the disease. We observed that DA-macroglia were also present throughout the parenchyma of the MS sample with active lesions but were highly enriched within the WM of the sample with inactive lesions, in which they were particularly abundant. ISS also unveiled novel sub-compartments within MS lesions, allowing improved resolution for human MS lesion classification when compared to classical neuropathology. With this study, we also demonstrate for the first time how cellularly resolved in situ transcriptomics can be applied to map cellular states and neighbourhoods in disease vs control studies and that such data can generate novel insights into disease processes and dynamics.

Our study reveals spatio-temporal cellular dynamics of neuropathology in EAE with some aspects likely translated to human MS and highlights the transitional plasticity of glia in response to inflammation. We provide a web resource where these spatial datasets can be explored, https://tissuumaps.scilifelab.se/2023_spinal_brain.html, which will constitute an important resource for the MS and neuroscience community to further investigate the function of the different cell populations involved in MS and their contribution to disease processes.

## RESULTS

### Comprehensive spatial and temporal cellular mapping of pathology in the EAE mouse model of MS

MS is characterised by multifocal lesions and covert pathology in normal-appearing white and grey matter regions. We first leveraged direct RNA-targeted ISS in the EAE mouse model of MS to probe early, acute and resolving inflammatory pathology that arises at different time points following disease induction. We then spatially characterised the cellular dynamics corresponding to these processes (**Figure 1A**). EAE was actively induced in C57BL/6J mice with murine MOG35–55 peptide, pertussis toxin (PTx), and complete Freund’s adjuvant (CFA); and PTx/CFA treated mice were used as controls (**Figure 1A, I.**). As expected, we observed the classical EAE disease course, with ascending flaccid paralysis that mirrors spinal cord lesion occurrence^36^ **(Supplementary Figure 1A**). To capture lesions at different stages of inflammation and to build a pseudotime of lesion development, we examined distinct CNS anatomical regions at the peak of EAE and at late stages. We collected coronal brain (≅Bregma 1.1) and transverse caudal-to-cranial spinal cord tissue sections (including cervical (C), thoracic (T), and lumbar (L) areas) from 4 representative control and 4 EAE mice of both sexes at peak (score ≥ 3, see Methods, **Figure 1A, I., Supplementary Figure 1A**). Additionally, lumbar spinal cord tissue sections were collected 30 days post-symptom onset (PSO) (score ≥ 2) from 3 independent EAE and 3 control replicates to capture changes in cellular composition and distribution occurring at late stages of the disease (**Figure 1A, I., Supplementary Figure 1A**).

**Figure 1.**
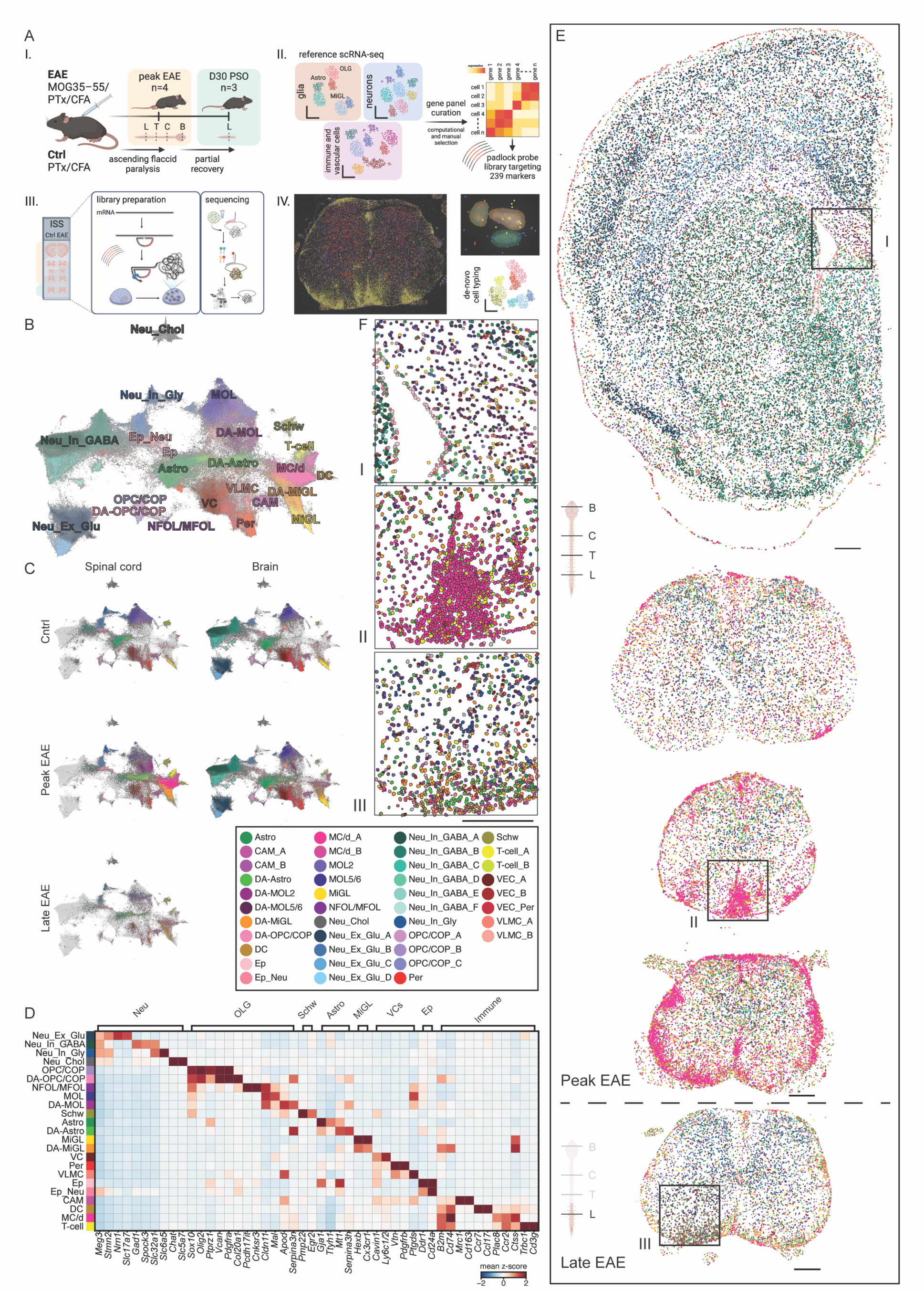
Comprehensive spatial mapping of homeostatic, disease-associated (DA), and immune populations across different anatomical regions of peak and late-stage EAE. (A) Experimental setup of the EAE induction and tissue collection, probe library design, ISS, and de novo cell typing. (I.) Induction of the EAE mouse model of MS and tissue collection time points. Three representative female and one male C57BL/6J mice from two independent EAE experiments were sacrificed together with their coupled controls at the chronic (peak) stage of the disease, and three representative female mice from two EAE experiments were sacrificed with their coupled controls at day 30 post-symptom onset (D30 PSO). Brains and spinal cords were isolated from the EAE and control mice. Brain (≅Bregma 1.1), cervical (C), thoracic (T), and lumbar (L) spinal cord (SC) tissue sections of 10μm thickness were collected for ISS of the peak, and L SC sections were collected for ISS of late stage EAE. (II.) A total of 239 genes were selected based on scRNAseq datasets and used for the padlock probe library preparation to target cell populations in their native positions. (III.) ISS workflow. (IV.) An example of a lumbar spinal cord tissue section with total reads generally corresponding to the immune cells (yellow), neurons (red), and glia (blue). Right is a zoom-in of three DAPI-segmented cells with corresponding reads and schematics of a UMAP depicting clusters annotated based on marker gene expression. (B) UMAP depicting cell populations defined by ISS and denovo cell typing of EAE and control SC and brain sections. Labels on the UMAP correspond to the 23 main annotated cell types, which were further subclustered at higher resolution into 41 clusters (see colour-coded legend in C). (C) Distributions of annotated cell populations between the spinal cord and brain sections from peak and late EAE. (D) A heatmap with selected differentially expressed genes across main cell types. Shown is the mean z-score. (E) Representative spatial maps of cells colour-coded by cluster type (see legend in C) across different CNS regions from the peak and late EAE stage. (F) Insets showing zoom-ins of selected regions from E, highlighting the cell types with immune infiltrates. Scale bars – 325μm. Astro - astrocytes, CAM - CNS-associated macrophages, MOL - mature oligodendrocytes, MiGL - microglia, OPC/COP - oligodendrocyte precursor cells and differentiation-committed precursors, DC - dendritic cells, Ep - ependymal cells, Ep_Neu - mixed ependymal and neuronal cells, MC/d - monocytes and monocyte-derived cells, NFOL/MFOL - newly-formed and myelin-forming oligodendrocytes, Neu_Chol - cholinergic neurons, Neu_Ex_Glu - glutamatergic excitatory neurons, Neu_In_GABA - gabaergic inhibitory neurons, Neu_In_Gly - glycinergic inhibitory neurons, Per - pericytes, Schw - Schwann cells, T-cell - T cells, VEC - vascular endothelial cells, VLMC - vascular leptomeningeal cells, VC - vascular cells including VEC and VEC_Per populations.

We anchored our ISS probe panel design in published single-cell transcriptomic datasets in this model^14, 22, 37, 38^ and focused on markers targeting classes of homeostatic and disease-associated (DA) glia, i.e., astrocytes, oligodendroglia, and microglia, as well as their tissue-neighbouring populations of neurons, vascular cells, and infiltrating immune cells (**Figure 1A, II.**). The final panel comprised 239 marker probes (**Figure 1A, II.**, **Supp. Table 1**, High Sensitivity CartaNA kit, now 10x Genomics^35^) that could be used in unique combinations to elucidate our targeted populations (**Figure 1A, II. - IV.**). ISS was performed on the intact EAE and control tissue sections as described previously^35^ (**Figure 1A, III.**). In brief, libraries were prepared, where target-specific padlock probes (PLPs) hybridised to mRNA. These PLPs were then ligated and amplified with rolling circle amplification (RCA) to generate rolling circle products (RCPs). The identity of each RCP was determined by a cyclic readout using fluorescently labelled probe constructs, and subsequent RCP decoding uncovered the vast expression landscape across the collected tissues (**Figure 1A, IV.**). Because of the increased sensitivity of the High Sensitivity CartaNA kit, we encountered issues with overlapping RCPs (optical crowding) in the 2D projected data we used for decoding. This was resolved by first deconvoluting the images of a subset of the samples using Deconwolf^39^, and using these deconvolved images as a training set for content-aware image restoration (CARE)^40^. The RCPs in the CARE processed 2D projected images were much sharper, leading to fewer overlapping RCPs. This resulted in an average 2.41±0.60 fold increase in the number of decoded transcripts. Next, nuclei-stained images were segmented to generate cellular masks, and the reads were assigned to individual cells as described in Methods (**Figure 1A, IV.**). Due to the varying density of cells across the tissue section, we improved the segmentation by training the existing 2D_versatile_fluo model in StarDist^41^ with manually segmented cells in the lesion areas. We noted an increase of 1.25±0.22 in the number of cells segmented with the improved model. Various available computational tools use the alignment of spatial data with scRNAseq to determine the identity of each cell based on the reads assigned^42–46^. Although several scRNAseq datasets were generated targeting specific populations in EAE^14, 18, 21, 22^, they do not incorporate in-depth molecular characterisation of all the populations of our interest. Therefore, we developed a novel hierarchical unsupervised clustering strategy with de-novo cell typing (**Figure 1A, IV.,** see Methods). By *de novo* clustering and cell type classification, we avoided the imputation of cellular identities and any other biases, such as sampling biases, that could have been introduced. This approach allowed us to characterise and spatially map single cells directly in the tissues and quantify their proportions.

### In situ sequencing resolves major cell populations and states in EAE

Upon ISS of all selected anatomical regions from control, peak, and late EAE samples, we recovered a total of 499,978 cells (across 304 mm^2^ of tissue); 178,435 from the spinal cord and 321,543 from brain sections, with an average cell containing 25±14 reads corresponding to 14±7 distinctive genes **(Figure 1B-C, Supplementary Figure 1B)**. Following hierarchical clustering, we annotated 41 unique populations, covering major homeostatic, disease-associated CNS-resident, and immune cell types (**Figure 1B-F, Supplementary Figure 1C-E**). Within the control spinal cord and brain samples, we spatially resolved all major CNS-resident homeostatic populations **(Figure 1B-D, Supplementary Figure 1C-D)**. Neurons (Neu; *Meg3+* and *Stmn2+*) represented the largest fraction of cells **(Figure 1D, Supplementary Figure 1E)** and were predominantly brain-derived or located within the grey matter (GM) of the spinal cords, as expected (**Supplementary Figure 1D, Supplementary Figure 2A)**. We could further subdivide them into excitatory (Ex) glutamatergic (Glu; *Nrn1+* and *Slc17a7+*), inhibitory (In) gabaergic (GABA; *Gad1+*, *Spock3+, Slc32a1+*), glycinergic (Gly) (*Slc32a1+* and *Slc6a5+*), and cholinergic (Chol; *Chat+* and *Slc5a7+*) types^38, 47–49^ **(Figure 1D, Supplementary Figure 1C)**. While some neuronal clusters were detectable throughout the brain, others were distributed within specific regions, such as the striatum, cortex, or specific cortical layers **(Supplementary Figure 2A)**.

Also highly abundant were oligodendroglia (OLG; *Sox10+*, *Olig2+, and Plp+*) **(Figure 1B-D, Supplementary Figure 1E,** https://tissuumaps.scilifelab.se/2023_spinal_brain.html), densely populating corpus callosum (CC), and white matter (WM) of the spinal cord, as expected **(Figure 1E, Supplementary Figure 1D, 2B)**. Main targeted OLG subclusters exemplified three major stages of lineage developmental trajectory, i.e., oligodendrocyte precursor cells and differentiation-committed precursors (OPC/COP), intermediate states of differentiation including newly-formed and myelin-forming OLG (NFOL/MFOL) and mature (MOL) subpopulations^37^. OPC/COPs were identified upon upregulation of *Olig2* and other progenitor markers, including *Ptprz, Vcan, and Pdgfra*^37^, while NFOL/MFOL expressed genes involved in early stages of differentiation, such as *Pcdh17it (9630013A20Rik)*^50^, *Mpzl1*^37, 51, 52^ and marker *Cnksr3*^37^ **(Figure 1D, Supplementary Figure 1C,** https://tissuumaps.scilifelab.se/2023_spinal_brain.html). Myelination-related genes (*Cldn11*, *Mal,* and *Mog*) were also expressed by NFOL/MFOL and further upregulated in MOL clusters. It was previously demonstrated that MOLs are transcriptionally heterogeneous, with unique subpopulations exhibiting spatial preferences for specific anatomical regions within the mouse CNS^53, 54^. Our analysis captured two major MOL populations, MOL2 and MOL5/6, based on the combination of markers, including *Klk6* and *Ptgds*, respectively **(Supplementary Figure 1C)**. In accordance with the literature, most of the brain-populating MOLs were of MOL5/6 identity, while within the spinal cord tissue, MOL2 were more abundant and enriched within the WM^37, 54^ **(Supplementary Figure 2B)**. Since parts of the dorsal root ganglia (DRGs) were preserved during the spinal cord isolation, we recovered a cluster of Schwann cells (Schw) that shared several markers with OLG but differentially expressed higher levels of *Pmp22* and *Egr2*^55^ **(Figure 1B-E, Supplementary Figure 1D, 2B)**. Schwann cells assigned to the spinal cord and brain regions are presumably artefacts arising due to our marker panel limitations. Nevertheless, we cannot exclude the possibility that some of these cells are indeed Schwann cells, especially in EAE^56^.

Expectedly, amongst highly prevalent populations in control tissue sections were also astrocytes (Astro; *Gja1+, Gpr37l1+, Gfap+,* and *Aqp4+)* and microglia (MiGL; *Hexb+, Aif1+, Cx3cr1+, Olfml3+,* and *Tmem119+*) **(Figure 1B-D, Supplementary Figure 1D-E, Supplementary Figure 2B,** https://tissuumaps.scilifelab.se/2023_spinal_brain.html). Finally, our probe panel allowed us to annotate a broad population of CNS-resident vascular cells (VCs) consisting of pericytes (Per), vascular endothelial cells (VEC), and vascular leptomeningeal cells (VLMCs)^55^. VECs^38^ were enriched for vascular genes, including *Ly6c1/2* (probe targeting homologous region of *Ly6c1* and *Ly6c2*), but did not express VLMC (*Pdgfra* and *Ptgds*) or Per (*Pdgfrb* and *Vtn*) markers **(Figure 1B-D, Supplementary Figure 1C)**. VLMCs were strongly labelling meninges of the brain and spinal cords, and VEC/Per cells were, in some cases, resembling vessel organisation **(Supplementary Figure 2C)**. Even though our sequencing panel did not include any classical markers for ependymal lineage cells, we observed a unique population of cells surrounding the ventricle and the central canal of the spinal cord, which we annotated as ependymal (Ep; *Ddr1+, Cd24a+*) or a mix of ependymal and neuronal (Ep_Neu; *Cd24a+, Meg3+*) cells **(Figure 1D, Supplementary Figure 2C)**. Finally, 18,072 cells (3,6% of total cells) were assigned a spectrum of canonical genes marking various cell types, and we could not define their identity. These cells were not limited to any particular areas and were consequently annotated as Unknown and excluded from the downstream analysis.

All major CNS-resident populations identified in control samples were retained in EAE, where disease-associated glial states and immune populations were prevalently detected **(Figure 1C, E-F)**. Most immune cells were infiltrated myeloid cells, distributed multifocally throughout the WM of the peak EAE spinal cords **(Figure 1C-F, Supplementary Figure 2D)**. We recovered myeloid subclusters upon their robust expression of genes related to antigen presentation, but also specific markers for CNS-associated macrophages (CAM; *Mrc1+, Cd163+*, and *Cbr2+*), dendritic cells (DC; *Ccl17+* and *Ccr7+*), and monocytes/monocyte-derived cells (MC/d; *Plac8+*, and *Ccr2+*) **(Figure 1D, Supplementary Figure 1D)**. Expectedly, T-cells (*Trbc1+, Cd3g+,* and *Cd4+*) were also EAE-specific, with the highest prevalence within the peak spinal cord samples **(Figure 1C-F, Supplementary Figure 2D**). We were able to resolve disease-associated glial states based on their expression of interferon (IFN) response genes, serine protease inhibitors (Serpins), and genes involved in major histocompatibility complex (MHC) I and II, among others **(Figure 1C-E, Supplementary Figure 1D,** https://tissuumaps.scilifelab.se/2023_spinal_brain.html). Some markers, such as *Col20a1*, *Apod*, and *Mt1*, were upregulated only in disease-associated populations of OPC/COPs, MOLs, and astrocytes, respectively, while MHCI and II genes were most prominently upregulated in disease-associated microglia, as previously reported^57^. Disease-associated glia were the most abundant within the spinal cords at peak EAE. Importantly, they were also detected within the brain and the late-stage spinal cords (**Figure 1C**), suggesting both early induction and persistence of DA-glia cell states along the EAE disease course.

### Immune populations within spinal cords at peak EAE diminish in a caudal-to-cranial manner but infiltrate the brain

Upon induction of EAE with MOG35-55, myelin-specific T cells progressively infiltrate the spinal cord in an ascending manner, which triggers multifocal lesion formation^36, 58^. However, the brain in this EAE model does not develop pronounced lesion pathology. We first spatially mapped and quantified the proportions of immune populations across the lumbar, thoracic, and cervical spinal cord and the brain **(Supplementary Figure 3A)**. Immune infiltrates were indeed diminished in a caudal-to-cranial manner along the spinal cord **(Figure 1E)**. For example, Monocytes/MC-derived cells (MC/d), but also CAMs, DCs, and T-cells, showed a significant decrease in densities from lumbar to cervical sections (**Supplementary Figure 3A,** p < 0.05, n = 4). We also observed that sporadic immune cells, including MC/d, DCs, and T-cells, began infiltrating the brain parenchyma **(Figure 1E, F, Supplementary Figure 1D, 2E)**. These immune populations were enriched at the medial side of the brain, near the ventricle and within the medial corpus callosum, but also at the ventral side of the brain proximal to the meninges, indicating two different locations of infiltration in the brain **(Supplementary Figure 1D, Supplementary Figure 2E, Supplementary Figure 3A, B).** Notably, the variability in immune infiltration rates across the four replicates suggests that we are capturing the earliest stages of brain inflammation (**Supplementary Figure 1D)**. These results suggest spatial preferences of lesion initiation in MS and are in accordance with previous findings that lateral ventricles and corpus callosum increase in volume at peak EAE^59^ and that the most noticeable immunopathological changes develop at the blood-brain-barriers (BBB) of parenchymal veins and at the ventricular sites^10, 60, 61^.

### Inflammatory dynamics differ between spinal cord anatomical regions and tracts

It has been shown that clinical subtypes of MS display different distributions of spinal cord lesions, with the dorsal column being frequently affected across subtypes and lateral funiculi being more often targeted in primary progressive and secondary progressive compared to relapsing-remitting patients^62^. We then investigated in further detail the dynamics of immune infiltration within different spinal cord areas. We observed that mild-to-moderate inflammation, as exemplified in peak EAE cervical cords, materialises as meningeal infiltration of immune cells throughout the superficial WM (**Figure 1E, Supplementary Figure 1D)**. This infiltration aggravates within the dorsal and ventral column at the site of the ventral medial fissure **(Figure 1E, Supplementary Figure 1D)** and potentiates further across ventrolateral sites, as seen in thoracic and lumbar EAE sections **(Figure 1 E, Supplementary Figure 1D, Supplementary Figure 3A-B, Supplementary Figure 4B**). We then examined how the immune load changes over time by comparing lumbar sections from the peak and late EAE. We observed a substantial decrease in the abundance of immune populations, including T cells, MC/d cells, and DCs, with time (**Figure 1E-F** and **Supplementary Figure 3B,** p < 0.05, n = 3), consistent with resolving inflammation during the late phase of the disease. Immune populations were particularly preserved at these late stages within the superficial WM at ventral sites but, unexpectedly, also within the dorsal column where the ascending tracts run (**Figure 1E, Supplementary Figure 4C**). The immune infiltrates remained abundant, specifically within the regions of gracile fasciculus and postsynaptic dorsal column pathway, across replicates. Similarly, the inflammation persisted within the contralateral regions where the descending rubrospinal tracts are localised (**Supplementary Figure 4C**). While certain anatomical regions appear to be more prone to develop lesions, our data also suggest that specific areas could be less efficient in resolving inflammation, which might also contribute to lesion chronicity. Taken together, we show that cross-regional mapping of CNS at peak and late stage EAE allows modelling of inflammation from initiation to acute and resolving stages and that certain anatomical regions and neuronal tracts might be more susceptible to lesion development and chronicity.

### Cell composition-driven identification of five distinct pathological compartments across regions and disease time points

EAE lesions are commonly identified based on increased nuclei density reflecting immune infiltration, immunostaining for lesion-enriched microglia, macrophages and/or loss of myelin^63, 64^. Such methods provide binary discrimination between the lesion and non-lesion tissue and introduce biases through thresholding for myelin or nuclei stains. We, therefore, investigated whether we could more precisely identify lesions or even minor pathological changes by analysing cellular neighbourhoods. We based our analysis on the 23 main specified cell types (**Figure 1B**) and unbiasedly probed the types of neighbourhoods that they form across the spinal cord and brain tissue sections (see Methods, **Figure 2A**). We examined the local neighbourhood around each cell, defined as a 100μm radius, which generated a neighbourhood matrix for each cell that we then clustered and annotated (**Figure 2A-C**, see Methods). Similar methods have been used previously to identify healthy neuroanatomical compartments based on spatial data^65–68^. In control spinal cords, the neighbourhood analysis allowed for annotating major anatomical regions, for instance, GM, WM, the central canal, and preserved DRGs **(Figure 2B-C, Supplementary Figure 5A**). In the control brain, we could also distinguish between the corpus callosum, cortex, compartment surrounding the meninges, the ventricular layer, striatum, and a compartment overlaying septal nuclei, hypothalamus, and substantia innominata (SN_HY_SI), confirming the power of our unbiased computational approach to identify compartments with diverse cellular compositions (**Supplementary Figure 5A)**.

**Figure 2.**
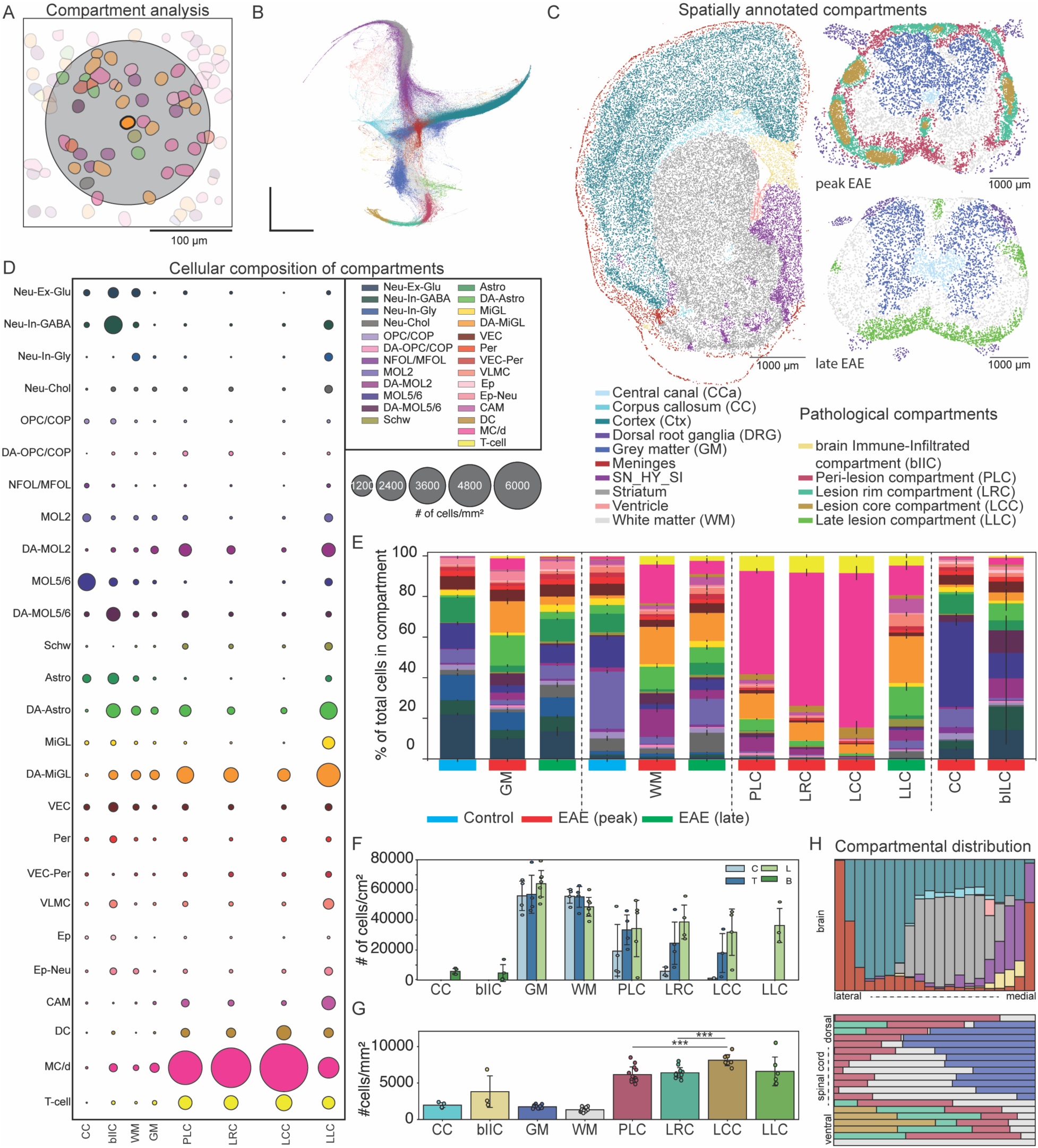
Distributions and cellular compositions of pathological compartments. **(A)** The automated compartment analysis. For each cell, neighbouring cells within 100μm were considered to be a part of its neighbourhood. The neighbourhood composition matrix was subsequently clustered and annotated. **(B)** The force-directed graph drawing of the neighbourhoods coloured by the annotated compartments (See legend in C). **(C)** Polygons of expanded nuclei coloured by the neighbourhood-driven compartment annotations, including disease-associated and anatomical compartments. **(D)** Dot plot with the numbers of cells per mm² across five disease-associated compartments. EAE corpus callosum (CC) and EAE WM and GM are depicted for comparison. **(E)** Stacked bar chart showing the relative cellular composition of the different anatomical and disease-associated compartments. **(F)** Bar plot showing the average number of cells per mm² defined as either one of the five disease-associated compartments or selected anatomical compartments from EAE. Error bars depict standard deviations. **(G)** Bar chart showing the average number of cells per mm² across the compartments defined in F. Error bars showing the standard deviations. **(H)** The lateral to medial gradients of the compartments within one representative brain section and dorsal to ventral gradients in one representative spinal cord section (see legend in C).

By applying the same approach to the EAE samples, we resolved equivalent anatomical regions but also additional compartments corresponding to lesions (**Figure 2C, Supplementary Figure 5A)**. Moreover, we identified distinct sub-compartments within lesions and disease-affected regions. Overall, we annotated five distinct disease-induced pathological compartment types; one corresponding to the immune-infiltrated callosal region in the brain, three being abundantly distributed throughout the white matter of the spinal cords at peak EAE and a fifth compartment only present in the lumbar spinal cord from late EAE stages (**Figure 2C)**.

### Disease-associated glia and vascular cells are hallmarks of the brain immune-infiltrated compartment (bIIC)

Even though brain lesions typically do not develop in this type of EAE, we identified pathological compartments predominantly within the medial corpus callosum at the superior side of the ventricle, which we annotated as the “brain Immune-Infiltrated compartment” (bIIC) **(Figure 2C, 3A, Supplementary Figure 5A)**. Although we detected these immune infiltrates within all four brain replicates at peak, only the two most affected sections exhibited clear bIICs **(Supplementary Figure 5A, C)**. Noteworthy, the neighbourhood composition of a bIIC was distinct from the pathological compartments within the spinal cords. It was overall similar to the corpus callosum but with increased cellular density (**Figure 2G**) and populated by immune infiltrates and disease-associated glial states (**Figure 2C-E, 3A)**. Lower abundances of homeostatic MOL types and increased numbers of DA-MOLs in this region suggest state transition of oligodendrocytes upon inflammation (**Figure 2D-E, 3A)**. This change, together with the overall lower numbers of MOLs in bIIC compared to the control callosal parenchyma, could explain previously reported myelination deficits and conduction abnormalities of EAE-affected corpus callosum^69^. On the other hand, homeostatic OPC/COP, Astro, and MiGL populations were preserved within the bIIC and complemented with their disease-associated counterparts **(Figure 2D)**. We also noted an increase in the number of vascular populations within the bIIC **(Figure 2D)**. This could reflect that the bIIC compartment was induced within a region with dense vasculature, which might act as a delivery pathway for the soluble cytokines to the brain parenchyma, promoting inflammation.

Taken together, we detect initial phases of brain pathology at peak EAE that are closely related to acquired disease-associated glial profiles in response to inflammation. Since brain lesions in this EAE model typically do not emerge, the transitions of bIIC compartments into mature lesions might thus require additional environmental triggers.

### Differential spatial distribution of spinal cord lesion compartments at peak EAE

The pathological compartments identified within peak EAE spinal cord differed in their cellular neighbourhoods but also in-tissue localisation (**Supplementary Figure 5)**. We annotated these compartments based on their distribution within lesions with fully developed, complex architecture (Lesion stage III in **Figure 3B-C**) as: 1) lesion core compartment (**LCC**), 2) lesion rim compartment (**LRC**), and 3) peri-lesion compartment (**PLC**) (**Figure 2C)**. The lesion core compartment (LCC) was defined as being present at the centre of complex lesions (**Figure 2C)** and had the highest abundances of MC/d cells and DCs (**Figure 2D, E)**. It was also significantly more populated compared to PLC and LRC (**Figure 2G**). The lesion rim compartment (LRC) was commonly found in complex lesions surrounding the lesion core and presented a lower number of MC/d cells and DCs, and, conversely, a higher proportion of DA microglia, DA-MOL2 and DA-astrocytes (**Figure 2D, E)**. Peri-lesion compartments covered the outmost complex lesion area. PLCs had the highest content of glia, the majority of which were disease-associated (**Figure 2D, E**). DA-MOL2 and DA-Astro were abundant in PLCs and gradually reduced in the lesion rim and even further in the lesion core (**Figure 2D, E**). DA-MiGL showed the same trend but remained relatively prominent also within the lesion core compartment, although other monocyte-derived phagocytes were significantly more abundant in this region (**Figure 2D, E**).

**Figure 3.**
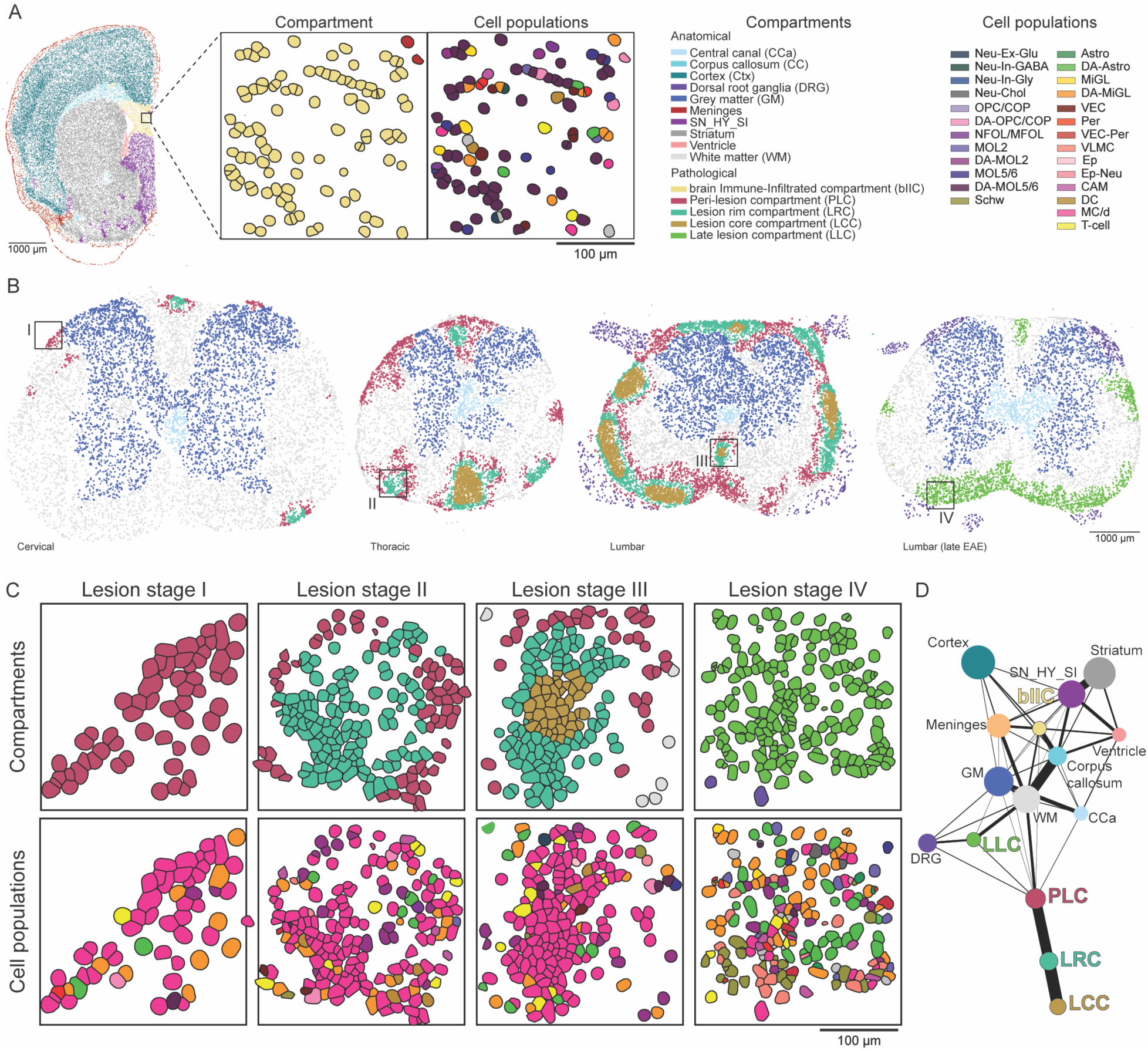
Active EAE lesions propagate in a centrifugal manner. **(A)** Polygons of expanded nuclei in one representative brain sample, coloured by the neighbourhood-driven compartment annotations. Inset shows a part of the bIIC and the corresponding cell populations. **(B)** Polygons of expanded nuclei coloured by the neighbourhood-driven compartment annotations, including disease-associated and anatomical compartments in representative spinal cords from the peak (cervical, thoracic and lumbar) and late (lumbar) EAE. **(C)** Insets showing the different lesion stages, corresponding compartments and cell populations. **(D)** Partition-based graph abstraction (PAGA) representation of connections between the annotated compartments. The size of the nodes represents the number of cells in the given compartment annotation, and the thickness of the edges represents the strength of the connection.

In accordance with observed differential inflammatory profiles across the caudal-cranial axis, peak lumbar spinal cord sections displayed the highest overall rate of disease-induced neighbourhoods, which were prominent across all four sequenced replicates (**Figure 2F, Supplementary Figure 5A)**. All thoracic sections contained peri-lesion compartments (PLC) and, to a lesser extent, lesion rim compartments (LRC), with three replicates also exhibiting some instances of lesion core compartments (LCC) (**Figure 2F, Supplementary Figure 5A)**. In cervical sections, only the presence of PLC was consistent across all replicates, suggesting that PLC is formed in response to the earliest pathological changes during lesion development (**Figure 2F, Supplementary Figure 5A).** Additionally, we observed an increase in the occurrence of disease-altered compartments in ventral regions, especially in the thoracic and lumbar sections (**Figure 2H, Supplementary Figure 5B**). Moreover, in peak lumbar sections, we noted alterations in the neighbourhood-based annotations of the posterior GM, where parts of the ventral horn were automatically assigned to the WM neighbourhood, suggesting certain similarities of this region with the WM **(Figure 2C, Supplementary Figure 5A)**.

### Centrifugal evolution of EAE spinal cord lesions

Longitudinal MRI has been previously used to investigate lesion development and evolution, with centrifugal and centripetal models being proposed^70^, despite MRI not allowing the probing of lesion composition at a cellular level. To investigate how complex active lesions develop in EAE, we analysed the compartmental dynamics in our spatial ISS data. Although bIIC was detected as a standalone compartment within the EAE-affected brain **(Figure 3A)**, we observed intricate spatio-temporal dynamics of different compartments within the EAE spinal cords **(Figure 3B)**. Fully developed lesions with complex architecture (Lesion stage III) were predominantly abundant in the acutely affected peak lumbar EAE spinal cord but absent in the cervical spinal cord (**Figure 3B)**. In contrast, we observed other lesions with simpler architectures composed only of one (Lesion stage I) or two compartments (Lesion stage II), primarily enriched in cervical and thoracic sections (**Figure 3B)**. Surprisingly, lesions with two compartments (Lesion stage II) comprised the lesion rim compartment (LRC) at their core, surrounded by the peri-lesion compartment (PLC) defining their outer edge (**Figure 3C).** These lesions were most prevalent in thoracic spinal cord sections and might thus constitute an intermediate stage during the process of lesion formation. We also found that the cervical and thoracic spinal cord presented an enrichment of lesions only consisting of one peri-lesion compartment (PLC) (Lesion stage I), indicating early-forming lesions (**Figure 3C)**. Thus, our data suggest that peri-lesion compartments (PLC) arise upon mild-to-moderate inflammation as standalone pathological compartments (Lesion stage I) and start spreading outwards while transitioning at their core into LRC and subsequently to LCC when inflammation progresses **(Figure 3B-C)**.

Next, we applied Partition-based graph abstraction (PAGA)^71^ to unbiasedly infer trajectories between the defined compartments (**Figure 3D**). We indeed observed transitions between the peri-lesion and lesion rim compartments, as well as the lesion rim and lesion core compartments (**Figure 3D**). Thus, our data indicate that active lesions in EAE are composed initially of sparse immune infiltrates and DA-glia. With increased immune infiltration and proliferation within the spinal cord, lesions compartmentalise and undergo a concentric centrifugal expansion, which nevertheless appears to be blocked when reaching the grey matter (**Figure 3B-C**).

### Resolved inflammation in late lesion compartments

While lesions at the peak EAE exhibit clear compartmentalisation, in late-EAE lumbar sections, the LCC, LRC, and PLC compartments are lost (**Figure 3C**). Instead, we observe a distinct type of disease-induced compartment of heterogeneous cellular composition, annotated as a fifth late lesion compartment (LLC). The LLCs are highly prevalent throughout the superficial white matter of the spinal cord tissue and predominantly located in ventral regions, suggesting that they are derived from resolving lesions (**Figure 3C, Supplementary Figure 5A-B**). The absence of other pathological compartments, including PLC, in these samples further supports the notion that there are no newly forming lesions at the late disease stage. GM in the late EAE samples was compositionally comparable to the control GM, and the late lesions were somewhat similar to the late-EAE WM with higher proportions of DA and immune populations (**Figure 2D, E)**. At the same time, the LLC is the compartment most populated with microglia and DA-astrocyte states, but also vascular populations and CAMs (**Figure 2D, E)**, which have been shown to reside in perivascular and meningeal structures^21^. This data suggests that lesions in late EAE are affected by gliosis, and the increase in vascular populations hints at a possible ongoing fibrotic scar formation, as previously reported^72^.

### Disease-associated glia mark global, lesion-independent pathology within the peak EAE spinal cords

Spinal cords at peak EAE exhibited not only focal compartments but also global alterations in cellular composition when compared to their CFA controls (**Figure 4A)**. These changes were noticeable already at the level of anatomical compartments. The GM and WM were populated by immune cells and disease-associated glia (**Figure 2D-E, 4A, B)**, consistent with the notion that the normal-appearing white and grey matter already exhibit disease signatures within microglia, astrocyte and oligodendroglia populations (**Figure 4D)**. Therefore, we proceeded with an in-depth exploration of DA glial transcriptional signatures and their spatial distributions.

**Figure 4.**
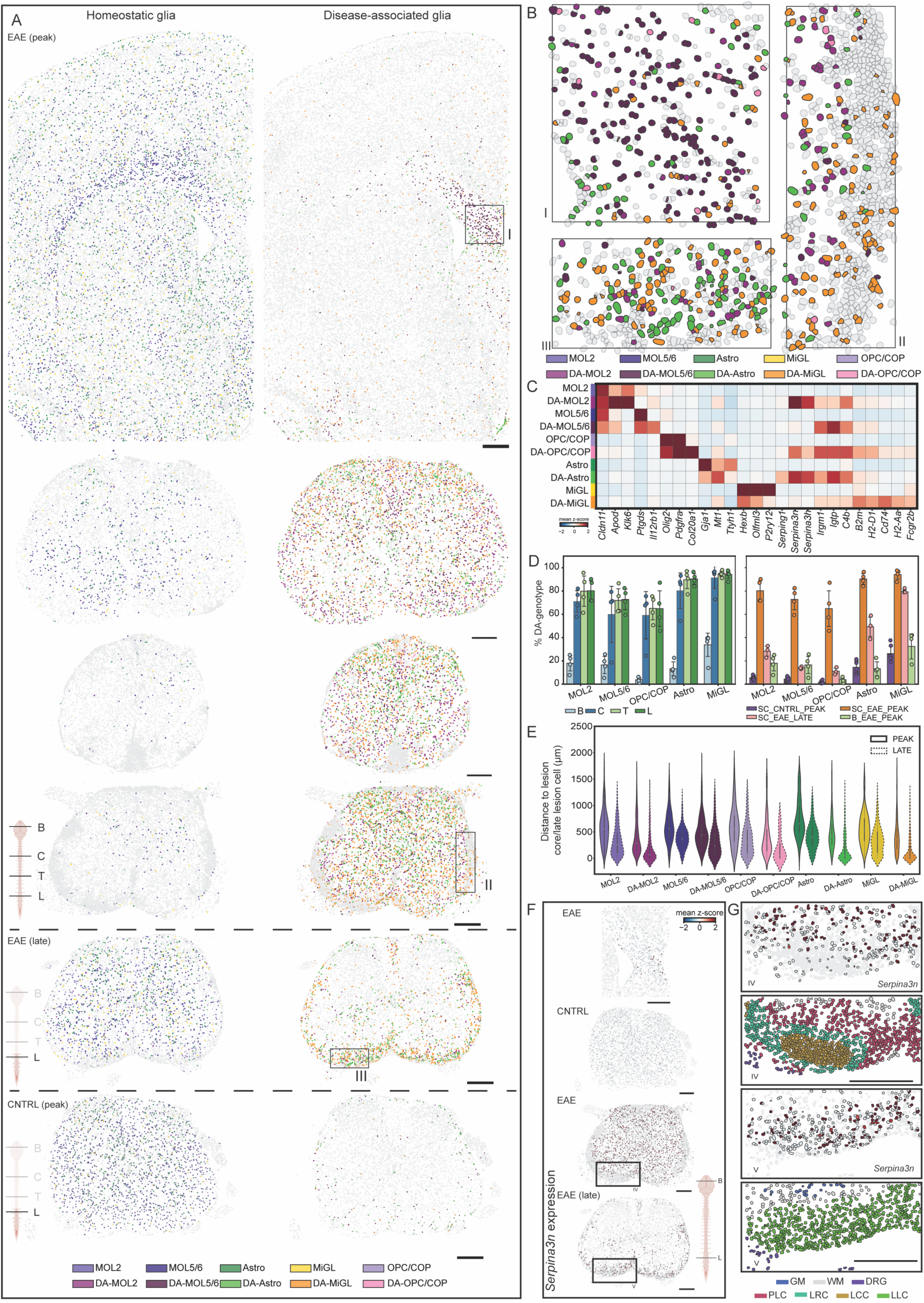
Spatio-temporal dynamics of homeostatic and disease-associated glial populations in EAE. **(A)** Spatial maps of homeostatic and disease-associated glial cells in the representative brain, cervical, thoracic, and lumbar spinal cord sections from peak EAE, a lumbar section from late EAE and a lumbar control section. **(B)** Insets showing zoom-ins of selected regions from A, highlighting the localisation of disease-associated glia. **(C)** Gene expression of selected marker genes. Shown is the mean z-score. **(D)** Rostral to cranial changes in the percentage of DA-genotype in glial populations (left) and temporal changes in the percentage of DA-genotype in glial populations (right). **(E)** Violin plots showing the distance to the nearest LCC defined cells, in the case of peak, and LLC defined cells for the late timepoint, for each of the (DA-) glial population. Violin plots showing data density, with median and 1.5 times the interquartile range. **(F)** Cells coloured by their expression of Serpina3n. Shown is the mean z-score. **(G)** Insets showing zoom-ins of selected regions from F, showing the cellular expression of Serpina3n in proximity to lesion regions. Shown is the mean z-score. Scale bars - 325µm.

The DA-MiGL exhibited decreased expression of homeostatic MiGL markers and upregulated membrane proteins related to their activation, including MHC genes, particularly *B2m* and *Cd74*, as reported previously in activated microglia (**Figure 4C**)^73^. Similarly, DA-Astros were recognised as reactive upon upregulation of *Gfap* and lower expression levels of classical homeostatic markers, including *Gja1* and *Ttyh1*. Moreover, they expressed high levels of Serpins (*Serpina3n, Serpina3h*), IFN response genes (*Igtp, Irgm1*), complement genes (*C4b, Serping1),* and metallothionein family genes *Mt-1* and *Mt-2*^74^ (**Figure 4C**). Within OLG, we recovered three unique DA clusters. DA OPC/COPs expressed *Col20a1*, Serpins, genes involved in IFN response, and antigen presentation via MHC I (*B2m, H2-D1)* and MHC II (**Figure 4C**). Similarly, DA MOLs expressed high levels of IFN and MHC-related genes, albeit to a lesser extent (**Figure 4C**). Notably, high expression levels of Serpins, but also *Apod,* were detected within DA-MOL2. In contrast, Serpins were not detected within DA-MOL5/6, which instead expressed *Il12rb1* and IFN-related genes (**Figure 4C**). Thus, markers involved in interferon response and antigen presentation were shared across all the DA glial classes, although their expression levels varied.

As mentioned previously, in the EAE brain, glial transitions into DA states were most prominent within the bIIC localised at the medial corpus callosum and upper ventricular area. DA-glia gradually decreased towards the lateral callosal site, which was, in turn, populated with homeostatic glial states (**Figure 4A, B, Supplementary Figure 6A).** Interestingly, microglia were broadly DA (up to 40%) also outside the immune-infiltrated hotspots, indicating global changes to the brain parenchyma at peak EAE (**Figure 4A, D, Supplementary Figure 6A)**. As opposed to the brain, DA-glia were remarkably prominent within peak EAE spinal cord sections and spread out throughout the parenchyma (**Figure 4A, D, Supplementary Figure 6B)**. In fact, most glia had acquired DA transcriptional profiles across cervical, thoracic, and lumbar sections, indicating global transitions not correlating with the lesion load (**Figure 4A, D).** Noteworthy, while the abundance of immune populations within lesion compartments increased with proximity to the lesion core compartment (LCC), homeostatic and disease-associated DA glial classes showed opposite trends, with DA glia most enriched within peri-lesion compartments (PLCs) compared to other pathological and anatomical regions (**Figure 2D-E**, **Figure 4A, B)**. This might be attributed to cell death and glial elimination within highly acute lesion core environments. Nevertheless, DA-MiGL were still relatively preserved also within the lesion core, which might be associated with their function in myelin clearance (**Figure 4A, B, Supplementary Figure 6B).** We examined how glial states change with increased distance from the lesion core and noted that homeostatic glial profiles are best preserved distal from lesions, mainly within the GM **(Figure 4E, Supplementary Figure 6B).** Also noteworthy, sparse DA-glia were found within the spinal cords of control mice treated with PTx/CFA (which triggers mild meningeal immune infiltration), predominantly within the WM (**Figure 4A**). This indicates that glia respond to inflammation, which is decoupled from lesion formation.

These results indicate that glia begin to acquire DA signatures in a localised manner upon early inflammation but eventually get globally affected, possibly due to the diffusion of soluble cytokines from lesions or through the broadly disrupted BBB. This data also supports the notion that the induction of disease-associated glial states and the formation of lesions, while interconnected, might occur independently, with the former possibly preceding the latter.

### Reestablishment of homeostatic and DA-oligodendroglia in late lesion compartments

Next, we studied whether the DA glial states persist along the disease course by comparing the spinal sections from the peak and late EAE. Strikingly, we noted a substantial reduction of DA states within oligodendroglia, including DA-MOL2, DA-MOL5/6, and DA-OPC in late EAE spinal cords (3 to 4-fold decrease compared to peak EAE) (**Figure 4D, Supplementary Figure 6B)**. Similarly, we observed a 50% decrease in the abundance of DA astrocytes from peak to late-stage EAE. Microglia, however, remained to a large extent in the DA state (80% of total microglia fraction) (**Figure 4D, Supplementary Figure 6B)**, indicating that neuroinflammation has long-lasting effects on microglia and, to some extent, astrocytes, while oligodendroglia seem to carry remarkable capacity for re-establishing homeostasis.

In contrast to their distribution at peak EAE, the DA glia at the late disease stage were found predominantly within the WM and lesion compartments (**Figure 4A, E, Supplementary Figure 6B, C)**. This could indicate the re-localisation of DA glia, especially migratory DA-MiGL, DA-Astros and DA-OPCs, to the lesion areas. Since myelinating oligodendroglia are presumably non-migratory, it is likely that DA-MOLs globally revert back to their homeostatic profiles upon decreased inflammation but are preserved within the areas where immune infiltrates remain present. This state transition was supported by transcriptional similarities between late EAE-derived homeostatic MOLs and DA-MOL states (**Supplementary Figure 7A-B).** Importantly, while MOLs were largely absent from lesions at peak EAE, they were re-established within the late lesion compartments (LLC), where the DA-MOL2 state was the most abundant **(Figure 2D**, **Figure 4A, B, Supplementary Figure 7C)**. Furthermore, we observed an upregulation of genes involved in lipid transport and myelination, including *Apod*, within oligodendrocytes in late lesion compartments (LLC) (**Supplementary Figure 7E)**. Taken together, our data suggest that MOLs in late lesion compartments (LLC) are newly generated, possibly involved in remyelination and that their regional identity is preserved during lesion recovery.

Nevertheless, not all glia within late lesions were DA. We noted that compared to peak EAE, where homeostatic glia were depleted from lesions, homeostatic glial states re-emerged in late lesion compartments **(Figure 2D, Supplementary Figure 6B, C)** and are found closer to the late lesion centre **(Figure 4E)**. Homeostatic oligodendroglia and astrocytes within late lesions may be newly generated or newly migrated and have yet to acquire disease-associated transcriptional signatures. One of the genes possibly involved in promoting the survival of these populations is *Serpina3n*, which encodes an inhibitor of the serine protease Granzyme B, involved in the cytotoxic function of T-cells^75^. *Serpina3n* was highly enriched in the parenchyma and peri-lesion compartments (PLC) at peak EAE and upregulated in the late lesion compartments (LLC) (**Figure 4F, G**). Since intravenous treatment of EAE mice with Serpina3n leads to neuroprotection and maintenance of myelin integrity^75^, *Serpina3n* expression might constitute a survival response by glial cells to tackle immune infiltration and avoid apoptosis, both at the initial stages and also at the late resolving stage.

### Disease-associated glia interaction hubs are preserved across all stages of EAE

Receptor-ligand pairs in single-cell transcriptomics datasets have been widely used to infer cell-cell interactions^76^. Nevertheless, these methods present several challenges^77^. Since the native position of each cell was retained in our ISS dataset, we could explore *bona fide* interactions between cells based on their proximity (**Figure 5A**). First, we computed the spatial neighbours’ graph by connecting the 5 nearest spatial components for each cell. This enabled us to both look at the number of connections formed between different cell types (**Figure 5C**) as well as an enrichment score (z-score), where the observed number of connections is compared to a randomly permuted set of connections (**Figure 5A, D,** see Methods).

**Figure 5.**
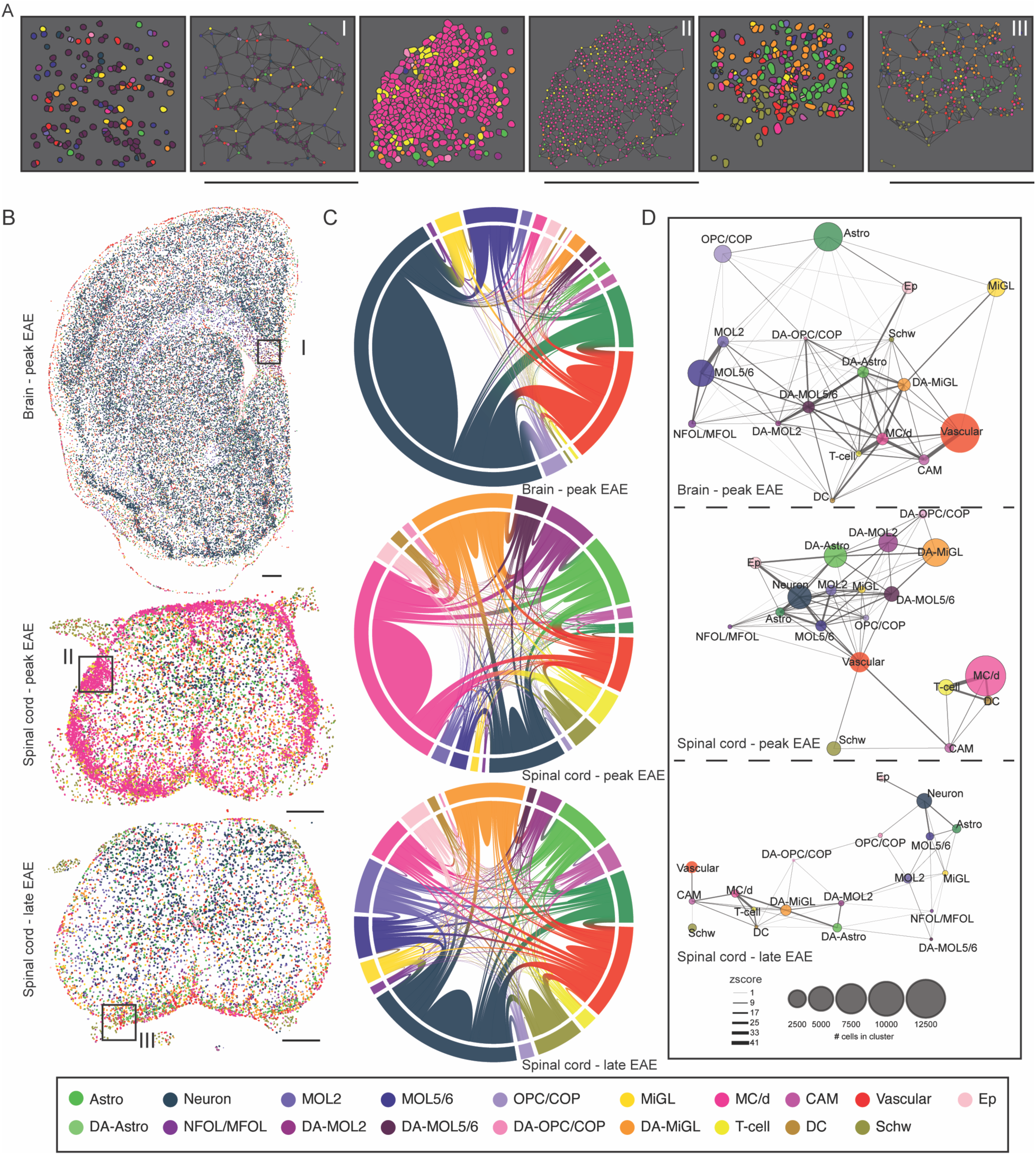
Disease-associated glia interaction hubs are preserved across all stages of EAE. **(A)** Insets showing zoom-ins from B, represented as polygons as well as dots with neighbour connections. **(B)** Polygons of expanded nuclei coloured by the cell type annotations. **(C)** Chord diagrams representing the numbers of connections between cell types from the spatial neighbourhood graph within peak EAE brain and peak and late EAE spinal cord. **(D)** A network representation of neighbourhood enrichment in EAE across the brain and spinal cord for peak and spinal cord for the late time point. The nodes represent the annotated cell types, with the size corresponding to the number of cells in the given cluster. The size of the edges corresponds to a significant enrichment (z score over 0). Scale bars - 325µm.

In the brain from peak EAE, we observed, as expected, that the most common interactions were between neurons and homeostatic glia, as well as neurons and vasculature (**Figure 5C**). However, when looking at the interactions occurring more often than random, we observed distinct hubs forming around immune infiltrates closely connected to the vasculature, DA-glia, homeostatic OLG, and a hub of OPC/COP-Astro-Ep connections, likely corresponding to the panglial syncytium^78^ (**Figure 5D**). Bridging immune and homeostatic hubs were DA glia that prominently interacted with each other but also with immune cells. For example, DA-MOL5/6 interacted mainly with DA Astrocytes, DA-microglia, but also T-cells and monocyte-derived cells (**Figure 5D**), which suggests that DA signatures in MOL5/6 might be induced either through direct or indirect signalling with these populations. Accordingly, the induction of a disease-associated immune state in oligodendroglia has been shown to be mediated, for instance, by IFN-gamma^14, 23, 79^, which is produced predominantly by T-cells^80^.

Within peak EAE spinal cords, the lesion-enriched immune cells formed a unique hub (**Figure 5D**). We observed a segregation of monocytes, dendritic cells and T-cells from other cell types, consistent with the formation of lesion core compartments (LCC) (**Figure 5D**). Notably, the DA-glia interacted more often with each other compared to their homeostatic counterparts. Disease-associated oligodendroglia, DA-MOL5/6 and DA-MOL2, retained at this stage a preferential interaction with DA-astrocytes and DA–microglia (**Figure 5D**), suggesting that the induction and/or maintenance of these cell states in the parenchyma, LRC and PLC might not be mediated by direct interactions with peripheral immune cells, in contrast to bIIC. In turn, neurons were surrounded by homeostatic astrocytes and oligodendrocyte lineage cells and, to a lesser extent, with DA-glia. Notably, DA-OPC/COPs displayed no prominent interactions with any homeostatic cell states, which could reflect loss of homeostatic functions as indicated by interactions of OPC/COPs with neurons and MOLs.

Within the late lumbar spinal cord, the immune hub was preserved, but especially MC/d cells were found to be prominently interacting with DA-MiGL, indicating their shared niche. The association of DA-oligodendroglia (DA-MOL2) with DA-astrocytes and DA-microglia was also present, which could suggest that newly generated MOLs might still acquire a DA state due to the presence of DA-astrocytes and DA-microglia upon resolving inflammation in LLCs. Notably, although both homeostatic and DA glia were present within close proximity in late lesions, they did not form prominent interactions with each other.

Thus, our data suggest that DA-glia are recurrent interaction hubs across regions and stages of EAE.

### Inference of ligand-receptor interactions by integration of spatial ISS with scRNA-seq

Having established *bona fide* spatial cell-cell interactions in EAE, we then leveraged two independent published scRNA-seq datasets of EAE at peak and control CFA samples^14, 22^ to infer ligand-receptor pairs that might play a role in the establishment, maintenance or functions of disease-associated glial states. We integrated the datasets to annotate the cell types following the ISS annotated nomenclature. To recover the most similar cell types, we used the top gene markers used in the ISS annotations as module scores combined with the originally published annotations. After curating the integrated dataset and discarding clusters with low numbers of cells, neurons and NFOLs, we inferred the cell-cell communications using the R package CellChat^81^.

DA-Microglia (but also homeostatic microglia) expressed several members of the CC chemokine ligands (CCLs) family that can bind to members of the CC-Chemokine Receptor (CCR) family (**Supplementary Figure 8)**. CCRs were, in turn, expressed in cell populations with which DA-microglia interact in the EAE brain (**Figure 5D**), such as T-cells and monocyte-derived cells, suggesting that these interactions could regulate cell migration towards the immune-infiltrated compartments, where DA-microglia were enriched **(Figure 5D, Supplementary Figure 6)**. In our spatial data, we observed interactions between DA-Microglia with DA-OPC/COPs in the brain, peak and late spinal cord (**Figure 5D**), and we found several interactions that could be regulating the OPC cell state. DA-Microglia express *Psap*, encoding for Prosaposin, which can bind in DA-OPC/COPs to *Gpr37l1*^82^ (**Supplementary Figure 8)**, encoding a glia-specific G protein-coupled receptor that has been shown to be necessary for OPC proliferation and differentiation^83^. Tnf-Tnfrsf1a signalling, which has also been shown to regulate OPC differentiation^84, 85^, was also suggested to occur between DA-Microglia and DA-OPCs/COPs (**Supplementary Figure 8)**. Furthermore, DA-Microglia were found to interact with DA-MOL5/6 and DA-MOL2 in EAE (**Figure 5D**) that, interestingly, express *Gpr37*, closely related to *Gpr37l1*, but instead inhibits oligodendrocyte differentiation and myelination^86^. Interactions involving Prosaposin and its receptors *Gpr37l1* and *Gpr37* were also found between DA-Astrocytes and DA-oligodendroglia in EAE (**Figure 5D, Supplementary Figure 8**). It has been recently shown that mature oligodendrocytes can participate in remyelination^87^, albeit in a non-efficient manner^88–90^. This could imply a reactivation of a myelination program in MOLs, which might be thus repressed by Gpr37 signalling triggered by Prosaposin from DA-microglia in the context of neuroinflammation. Thus, DA-Microglia and DA-Astrocytes in EAE might play an important role in regulating oligodendroglial cell states via, for instance, Prosaposin and TNF signalling.

We then focused on the receptor-ligand interactions of DA-MOL5/6 and DA-MOL2. As expected, both mature DA-oligodendroglial populations exhibited several pairs with both homeostatic and other DA-oligodendroglia (**Supplementary Figure 9**). For instance, DA-MOL secrete Pdgfa that binds its receptor Pdgfra in OPCs, which leads to cell proliferation (**Supplementary Figure 9**). We found ligand-receptor interactions between DA-MOL2 *App,* complement *C3/Jam3, C3ar1*/*Csf1 and Pros1* with DA-microglial *Cd74*, *Itgam/Itgb2*, *Csf1r* and *Axl*, respectively. DA-MOL5/6 presented similar interactions but also unique such as *Gas6* with DA-microglial *Axl* or *Mertkl* (**Supplementary Figure 9**). Thus, DA-microglia are also modulated by DA-oligodendroglial interactors, and some of these receptor-ligand interactions might be involved in the cross-induction or maintenance of these DA-glia cellular states.

### Spatial cellular mapping of human MS spinal cords reveals cytopathology of active and inactive lesions and spatial preferences among oligodendrocytes and astrocytes

To correlate our findings with spinal cord pathology in MS patients, we applied ISS with the *Xenium In Situ* platform to two human cervical spinal cord cross-sections bearing pathologically annotated active or inactive lesions^91^ (**Figure 6A**, see Methods). The active lesions were neuropathologically sub-divided based on HLA/PLP double staining into reactive (1; groups of HLA-positive microglial cells, no demyelination) and active lesions with HLA-positive ramified microglia (2.1) or intermediate (ramified-to-foamy) microglia (2.2). Inactive lesions were identified as (4) gliotic inactive hypocellular lesions with some HLA-positive microgliosis^91^. We employed the Xenium Human Brain Gene Expression panel with 266 genes targeting cell types and states implicated in physiological and pathological conditions (**see Methods**). The cells expressed, on average, 144±95 transcripts and 49±18 distinctive genes. In total, we identified 240,372 cells across two patient samples. We first confirmed the lesion co-localization based on MOG and HLA-DMB expression. HLA-DMB expression was enriched within regions annotated as active lesions where MOG expression was also detected (**Figure 6B**), but the levels varied depending on the lesion subtype, which generally correlated with PLP immunostaining. In contrast, inactive lesions were depleted from MOG, which was instead present within the WM and areas surrounding the lesions (**Figure 6B**). HLA-DMB expression was, expectedly, globally downregulated in the sample with inactive lesions when compared to the sample with active lesions and was detectable mostly within the WM and regions around lesion cores (**Figure 6B**). Importantly, we also identified a region contralateral to one of the inactive lesions that exhibited lower *MOG* expression (**Figure 6B**, marked with an asterisk). This region presented higher MOG immunoreactivity than the defined inactive lesions but slightly lower immunoreactivity when compared to surrounding WM **(Figure 6A)** and had thus not been classified neuropathologically as an inactive lesion.

**Figure 6.**
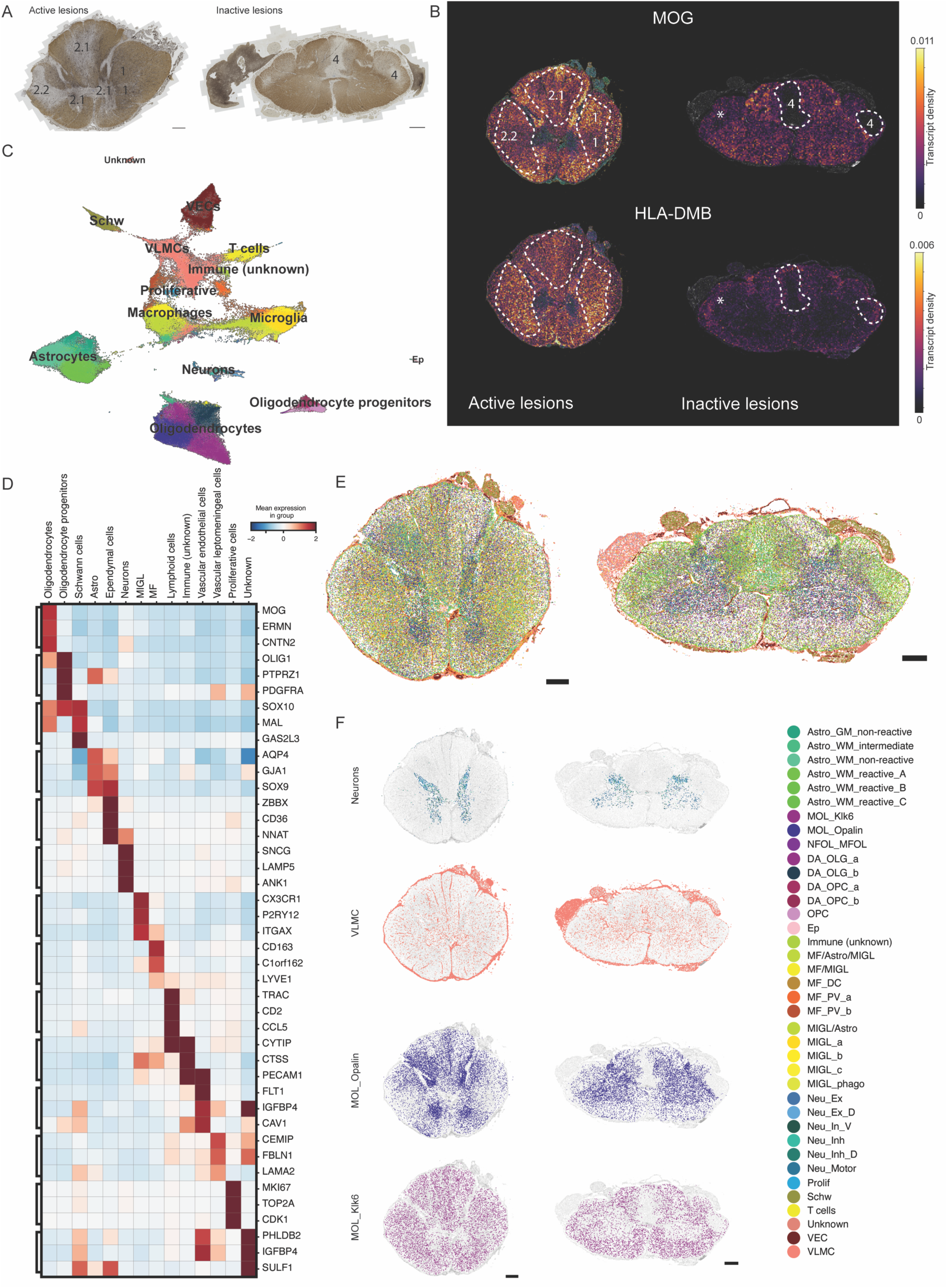
Spatial cell maps of human cervical spinal cord MS samples with active and inactive lesions. **(A)** Cervical cross sections from two MS patients with active and inactive lesions, pathologically annotated based on HLA/PLP staining. 1 - reactive lesions; 2.1 - active lesions with HLA-positive ramified microglia; 2.2 - active lesions with intermediate (ramified-to-foamy) microglia.; 4 inactive lesions; source and nomenclature: The Netherlands Brain Bank. Scale bars - 1000 ⲙm. **(B)** Cervical cross sections from mirror blocks (https://www.brainbank.nl/nbb-ms/characterization-of-ms-lesions/) of samples in A, used for Xenium In situ sequencing, depicting binned expression of MOG and HLA-DMB transcript density. Dashed lines outline regions with upregulated HLA-DMB expression (active lesions) and downregulated MOG (inactive lesions). Numbers correspond to pathological lesion staging. The asterisks mark a MOG-depleted area, pathologically not annotated as a lesion. **(C)** UMAP of cell types annotated using hierarchical clustering upon ISS. **(D)** A heatmap with selected differentially expressed genes across main cell types. Shown is the mean z-score. **(E)** Spatial maps of cell types. The colours of the cell types correspond to the colours in C. **(F)** Spatial maps of selected populations, including Neurons, VLMCs, MOL_Opalin, and MOL_Klk6. Scalebar corresponds to 1000µm.

Following ISS and cell typing with hierarchical clustering, we identified 38 cell populations covering major glial (astrocytes, oligodendrocyte lineage cells, microglia and Schwann cells), immune (macrophages, lymphocytes), neuronal, vascular and ependymal cell types (**Figure 6C-E, Supplementary Figure 10**). Some cell types exhibited expected spatial preferences, including GM-enriched neurons and vascular leptomeningeal cells (**Figure 6F)**. Moreover, we identified specific astrocyte and oligodendrocyte populations that were either WM- or GM-enriched (**Figure 6F, Supplementary Figure 10C**). This subdivision of human astrocytes agrees with previous findings^92^. The mature mouse oligodendrocyte populations MOL2 and MOL5/6 have been shown to preferentially reside within mouse spinal cord WM and GM, respectively^53, 54^. Here we demonstrate that classes of oligodendrocytes also exhibit spatial preferences within the human cervical spinal cord (**Figure 6F)**. Similar to the mouse MOL2, WM-enriched oligodendrocytes in humans expressed higher levels of *KLK6* but also *PCSK6* (**Supplementary Figure 10E)**. In contrast, GM oligodendrocytes were specifically labelled with *OPALIN*, previously identified as a marker of a distinct oligodendrocyte state within the human brain. This data suggests that GM- and WM-specific environmental cues induce distinct human MOL states or that neurons within these regions require specified oligodendrocyte support.

### Deconvolution of active human MS lesions into novel sub-compartments

Next, we utilised our automated compartment annotation framework using 24 main annotated cell clusters (**Figure 2A-B**) to identify anatomical and lesion compartments and pathological changes within normal-appearing parenchyma. We were able to annotate known anatomical structures, including the central canal (CCa), dorsal root ganglia (DRG), GM, and WM **(Figure 7A-B)**. We also resolved a vascular compartment (“vascular_a”), predominantly populated with vascular endothelial and vascular leptomeningeal cells and labelling the meninges with occasional parenchymal vessels in both sections **(Figure 7A-B)**. In addition, we could annotate four other vascular compartments populated with different degrees of immune infiltrates (“vascular_T-cell”, “T-cell_vascular”, “vascular_inf”, and “vascular_b”). **“Vascular_T-cell”** compartment co-localized with certain meningeal areas, especially in the active sample **(Figure 7A)**, and was mainly composed of VLMCs, but also T-cells and proliferative cells (population of cells expressing high degrees of proliferative markers including Mki67) (**Supplementary Figure 11B-C)**. These characteristics could indicate T-cell meningeal infiltration and that parts of this compartment might be capturing the meningeal follicle structures. The presence of Schwann cells within meningeal compartments (vascular_T-cell, vascular_a) is likely an indication that some DRG vasculature was captured within these areas. The **“T-cell_vascular”** compartment had, in turn, proportionally the highest levels of T-cells among all identified compartments (**Supplementary Figure 11B-C)** and clearly labelled parenchymal vascular structures within the sample with inactive lesions **(Figure 7B)**. Regions annotated as **“vascular_inf**” compartment also spatially resembled parenchymal vascular organisation in the sample with active lesions and were populated by vascular cells, immune infiltrates but also glial cells (**Figure 7A, Supplementary Figure 11B-C)**. While the “**vascular_b**” compartment contained vascular populations, it was enriched with a population of cells for which we could not determine a clear identity (“unknown” cluster) and limited to a small, possibly vascular structure in the sample with inactive lesions **(Figure 7B)**.

**Figure 7.**
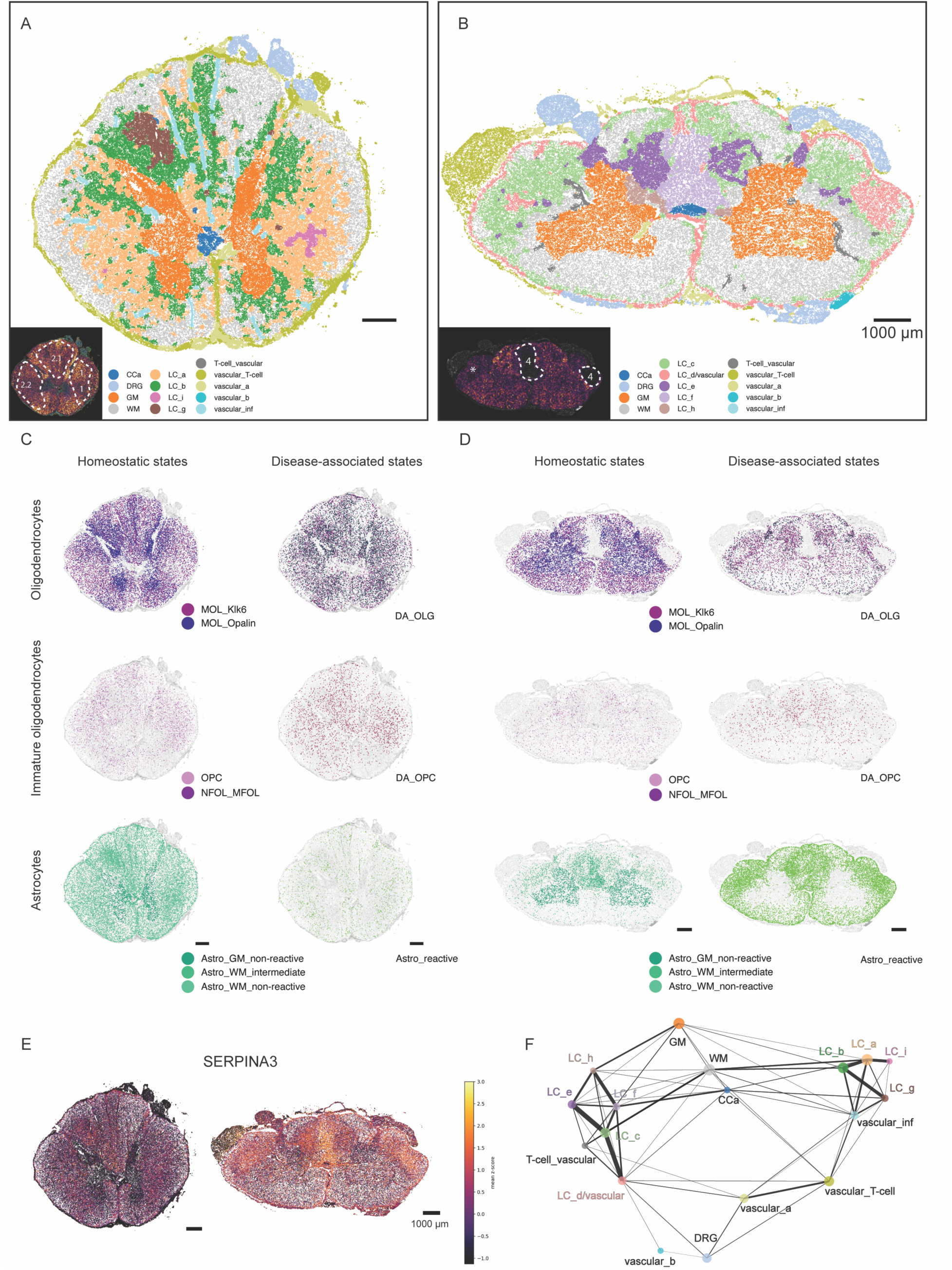
ISS unveils new lesion compartments and sub-compartments in human cervical spinal cord MS samples. **(A-B)** Identified anatomical and pathological compartments within the sample with active **(A)** and inactive **(B)** lesions. The same neighbourhood analysis approach was used as for the mouse ISS data. Binned MOG-expression maps (see Figure 6B) are added as a reference. **(C)** Spatial cell maps of the sample with active lesions depicting distributions of homeostatic and disease-associated macroglial states. Scale bars - 1000μm. **(D)** Spatial cell maps of the sample with inactive lesions depicting distributions of homeostatic and disease-associated macroglial states. Scale bars - 1000μm. **(E)** Spatial maps of the sample with active and inactive lesions with plotted SERPINA3 expression. Scale bars - 1000μm. **(F)** PAGA representation of connections between the annotated compartments. The size of the nodes represents the number of cells in the given compartment annotation, and the thickness of the edges represents the strength of the connection.

Importantly, we were able to identify nine compositionally and spatially distinct compartments corresponding to active/inactive lesions or disease-affected areas within the WM, which we defined as Lesions Compartments (**LC_a-i**) **(Figure 7A-B)**. These LC regions were present at the same anatomical locations as the four neuropathologically defined lesions (1, 2.1, 2.2 and 4, **Figure 6A**). For instance, regions annotated as LC_a, LC_b, LC_i, and LC_g were defined by cytopathological changes within the sample with active lesions, while LC_c, LC_d/vascular, LC_e, LC_f, LC_h marked disease-affected areas within the section with inactive lesions **(Figure 7A-B, Supplementary Figure 11B-C)**. Nevertheless, while we could identify WM lesions defined as 2.1, we were unable to detect the 2.1 GM lesions in the active lesion sample. It is possible that these lesions were present in the section assessed by staining but not in the sequenced sample (**Figure 6A-B**). Alternatively, we were not able to resolve them due to the lack of HLA-DMB (**Figure 6B**) and HLA-DQA1 (not shown) expression within these regions, and additional HLA markers could aid in their annotation.

In the active lesion sample, LC_a and LC_b were spread throughout the WM, with LC_b often adjacent to GM **(Figure 7A)**. When studying specific pathologically annotated active lesions in the context of our defined compartments, we noted that the WM “2.1” lesion (active lesions with ramified microglia) was predominantly composed of LC_b, which was indeed enriched with microglia compared to the WM **(Figure 7A, Supplementary Figure 11B-C)**, and marked with HLA-DMB expression **(Figure 6B)**. Interestingly, LC_b was also enriched with MOL_Opalin (**Supplementary Figure 11B-C)**, which, together with increased OPALIN and MOG expression within this compartment **(Figure 6B, F),** hints at potential ongoing remyelination and lesion repair. Elongated Vascular-inf compartments were also prominent in the “2.1” lesion, suggesting active neuroinflammation localised at vessels (**Figure 7A)**. The area affected by lesions annotated as “1” was, expectedly, marked with high HLA expression **(Figure 6B)** and broadly annotated as LC_a, which had proportionally higher numbers of microglia compared to the WM **(Figure 7A, Supplementary Figure 11B-C).**

While LC_a and LC_b presented a broad distribution, LC_g and LC_i compartments were focally distributed and marked restricted areas within WM lesions “2.1” and “1” (reactive lesions), respectively (**Figures 6A, 7A**). LC_g was distinct from surrounding LC_b mainly due to increased astrocytic abundance (**Supplementary Figure 11B-C)**. Noteworthy, while neuropathologically, the LC_a lesion was annotated as non-demyelinating, we were able to resolve a limited compartment, LC_i, characterised by a lower abundance of oligodendrocytes (**Supplementary Figure 11B-C).** Moreover, within this region, we detected lower levels of MOG **(Figure 6B)** and other myelin markers, including MAG, MOBP, and CLDN11 (see https://tissuumaps.scilifelab.se/2023_spinal_brain.html), suggesting pathological changes within oligodendrocytes in this area.

The annotated “2.2” lesion was compartmentalised in a similar way to its contralateral “1” lesion, despite the difference in demyelination **(Figure 6A**, **Figure 7A).** This could be explained by regional differences of pathologically and ISS-assessed sections or that myelin is affected in this lesion, but this was not reflected in oligodendrocyte abundance. In support of the latter, we do observe a prominent decrease in myelin genes, including MOG **(Figure 6B),** in this area.

### Enhanced power of ISS to unveil lesion compartments and sub-compartments not identified by neuropathology

The sample with inactive lesions did not display widespread pathological compartments but rather focal compartmentalisation, highly coherent with the pathological evaluation **(Figure 6A**, **Figure 7B)**. Moreover, as already suggested by *MOG* expression (**Figure 6B**), we could resolve sub-compartments of the annotated inactive lesions (**Figure 7B)**. While only one lateral lesion was histopathologically annotated in this sample **(Figure 6A)**, our compartment analysis revealed a similar compartmental composition of both contralateral sites **(Figure 6B** - lateral lesion “4” and the area marked with an asterisk), with a larger core compartment region within the annotated lesion **(Figure 6A**, **Figure 7B).** The lesion core regions were defined by the LC_d/vascular compartment, enriched with reactive astrocytes, vascular cells and perivascular macrophages. Interestingly, this compartment was also found to be labelling the meninges (**Figure 7B)**. The LC_c surrounding the lesion core compartments was spread throughout the WM and had a similar composition to the LC_d/vascular compartment but displayed a decreased vascular abundance **(Figure 7B, Supplementary Figure 11B-C)**.

In contrast to lateral lesions, the dorsal inactive lesion was mainly composed of LC_e and LC_f, surrounded by LC_c. LC_f was, in this instance, marking lesion core and was again enriched with vascular cells when compared to LC_e, but to a lower extent compared to the LC_d/vascular compartment (**Figure 7B, Supplementary Figure 11B-C)**. We also noted an adjacent GM area affected by the LC_h compartment that exhibited lower rates of astrocytes compared to the WM-affected areas in this lesion (**Figure 7A, Supplementary Figure 11B-C).**

### The newly identified lesion and pathological sub-compartments highlight the power of ISS to identify pathological changes not uncovered by histopathological analysis

To infer trajectories between the different compartments, we applied PAGA, as we had previously done with the EAE compartments (**Figure 3D**). We found three main trajectories emerging from WM (**Figure 7F)**. One trajectory linked to the inactive compartment LC_c, which then connected to LC_e and LC_d/vascular (**Figure 7F)**. The second trajectory connected WM to either the active compartment LC_a (which then linked to LC_i), and the third trajectory linked WM to the active compartment LC_b (which then linked to LC_g) (**Figure 7F)**. The second active trajectory was prominent in the dorsal WM, while the third trajectory was more ventral, which could indicate region specificity for lesion evolution into these sub-compartments.

### Distribution of disease-associated macroglia across human MS spinal cords is similar to the one in the mouse EAE model

Next, we studied the abundances and distributions of homeostatic and disease-associated (DA) macroglial states across the two human MS spinal cord samples. The DA-oligodendroglia and DA-astrocytes were found to be widespread throughout the samples with active lesions **(Figure 7C)**, consistent with our observation in the EAE model (**Figure 4**). In contrast, DA-astrocytes were limited to the WM of the inactive sample and further enriched within lesions **(Figure 7D)**, indicating astrogliosis. Their distribution was consistent with SERPINA3 expression **(Figure 7C-E)**, an orthologue of mouse *Serpina3n,* which was found upregulated in mouse late EAE lesions **(Figure 4F)**. While oligodendrocytes are depleted from inactive demyelinated lesions, the DA-oligodendrocytes were accumulated around the lesion cores and enriched within the LC_c, LC_e and LC_h but not in LC_f and LCd/vascular. Thus, our results indicate that several aspects of MS neuropathology, including certain glial states and their distributions, are common to the mouse EAE and human MS pathologies.

## DISCUSSION

In this study, we combine *de novo* cell typing by single-cell resolution spatial transcriptomics (ISS) with cellular neighbourhood analysis to investigate the spatial and temporal dynamics of disease-altered CNS regions in the EAE mouse model of MS and human cervical spinal cord MS tissue. By studying cellular neighbourhoods, we found five distinct lesion compartments in EAE and nine in human MS tissue, with profound differences in cellular composition and regional preferences. EAE compartments evolve from immune infiltrates through standalone-compartment lesions to complex architecture lesions consisting of three compartments, which resolve at later stages to distinct, non-compartmentalised lesions. We also found that the emergence of disease-associated (DA) microglia, astrocytes, and oligodendroglia occurs locally upon early inflammation. Eventually, glia get globally affected when inflammation becomes stronger, and most microglia, oligodendrocytes, and astrocytes acquire DA transcriptional profiles throughout the spinal cord parenchyma, independently of the lesion load. Thus, DA-glia induction might occur independently of lesions and possibly preceding their formation. To our surprise, at the late stage of EAE, most oligodendroglia and a large portion of astrocytes transitioned back to their homeostatic state while the DA profiles persisted within about 80% of microglia. Applying ISS to the human MS archival spinal cord tissue also provided a better understanding of human oligodendrocyte heterogeneity and the current understanding of human active and inactive lesion classification, with increased resolution and deconvolution into sub-compartments of distinct cellular compositions, and identification of new lesion areas not unveiled by classical neuropathological evaluation.

EAE is characterised by a caudal-to-cranial pattern of disease progression. In agreement with previous work evaluating inflammatory and other pathological changes in EAE with selected histological markers^63, 93^, tomography^94^, and MRI^59, 95^, we demonstrate that the brain from peak EAE can be utilised to exemplify mild inflammation and early tissue pathology. Rostral spinal cord regions can model lesion initiation and early lesion states, caudal spinal cord for acute lesion pathology and spinal cords from late EAE to epitomise resolving lesion environments. Indeed, we observed at peak stages a higher number of lesions with complex architecture present at lumbar spinal cord regions (**Figure 1-3**). These complex lesions are composed of three compartments (peri-lesion (PLC), lesion rim (LRC) and lesion core (LCC) compartments), and they were less prevalent in cranial regions, where lesions with one or two compartments (PLC and RLC) were more abundant. Therefore, given the distribution of these different types of lesions in a caudal-cranial gradient, our data led to a model in which active lesion formation in MS involves the following steps (**Figure 3B-C**): **1)** immune cell infiltration, **2)** formation of a “peri-lesion” compartment (PLC) with a focal concentration of peripheral immune cells and DA-MiGL (Lesion Stage I), **3)** development of a dual-composite lesion, where the “peri-lesion” compartment (PLC) at its core transitions into the “lesion rim” compartment (LRC; containing DA-glia, but in lower proportions when compared to immune cells) (Lesion Stage II), **4)** development of a multi-composite complex architecture lesion where the lesion core compartment (LCC), densely aggregated by mainly immune cells, is formed at the centre and is surrounded by the lesion rim compartment (LRC), and peri-lesion compartment (PLC) at the outer edge (Lesion Stage III), **5)** Finally, the inflammatory resolution leads to the formation of late lesions, primarily composed of DA-glia (Lesion Stage IV), but different from LRC.

Paramagnetic lesion rim has been previously described as a predictor of poor MS progression outcome^70^ and has been since recapitulated in the marmoset EAE model, where iron accumulated specifically within microglia/macrophages at the rim of older lesions^96^. The identified peri-lesion and lesion rim compartments in our EAE model emerged in early developing lesions at their core and re-localised subsequently to the border of fully developed active lesions. Hence, our data suggest that early pathological cellular neighbourhoods might be compositionally similar to the paramagnetic lesion rim but cannot be captured by MRI due to the absence of iron. What triggers the iron deposition at the late stages within these neighbourhoods remains to be determined^97^. It would also be of interest to investigate whether these neighbourhoods carry any other biomarkers, such as accumulation of DA-glia, that could be detectable at earlier pathological stages.

Different parts of the CNS behave as unique microenvironments with diverse susceptibilities for immune infiltration and, possibly, distinct cellular responses to inflammation^98^. Accordingly, our data indicates region specificity in the induction of EAE lesions **(Figure 1)**. In the brain, we observe immune infiltrates, including T-cells in the medial corpus callosum and ventricular zone, suggesting that these immune cells reach the brain with cerebrospinal fluid infiltrating the ventricles. We also observe immune infiltrates within and adjacent to the meninges, especially at ventral locations, suggesting an alternative delivery pathway. In the cervical mouse EAE spinal cords, the inflammation did not exhibit a clear dorsal-ventral gradient. Similarly, the formation of the Stage I lesions did not show a clear dorsal-ventral enrichment. Nevertheless, Stage II and III lesions were more abundant in ventral locations and dorsal column, which was consistent with increased inflammation within these regions of thoracic and lumbar sections (**Figure 1E**, **Figure 2H**, **Figure 3**). Inflammation persisted throughout the late EAE stage at specific locations, including superficial ventral WM, the dorsal column where ascending tracts are located, as well as contralateral regions where the descending rubrospinal tracts run. These preferences agreed with the locations of Stage IV lesions (**Figure 1E**, **Figure 3**). We demonstrate that the spatial occurrence of EAE lesions is hardly random and is, to a large extent, coherent with cervical lesions in MS patients^62^. Noteworthy, although our approach allowed the detection of covert pathological changes beyond lesion formation, we did not observe any such changes within the GM of the EAE model. Accordingly, GM might bear inhibitory signals that prevent the expansion of lesions throughout its parenchyma. It is also possible that WM-enriched MOL2 are more prone to get targeted by the disease, which leads to demyelinated WM lesions, while GM-enriched MOL5/6 might carry immunoprotective mechanisms.

Glial cell populations have been shown to acquire disease-associated states in EAE and MS^11–15, 20–22^. These DA states are characterised by the expression of genes involved in neuroprotection and immunomodulation, including interferon response genes. Thus, immune cells, for instance interferon-gamma producing T-cells, are candidates to induce these cellular states^14, 23–25^. We indeed observe prominent interactions of T-cells and DA-glia within the EAE brain (**Figure 5)**, which suggests that T-cells might play a role in the induction of glial DA states during early, localised inflammation. Nevertheless, these interactions were not captured in any other assessed spinal cord regions (**Figure 5**). Moreover, DA-glia were not more prevalent in lesion compartments where T-cells were preferentially residing, as for instance, within LRC and LCC. Namely, we noted a principally homogeneous distribution of DA-glia throughout the spinal cord parenchyma at peak, with certain enrichment of these states within PLC. Thus, alternative and possibly contact-independent mechanisms might be involved in their induction. Increased levels of cytokines have been detected in the serum, and cerebrospinal fluid of MS patients^10^, and immune infiltrates are considered to modulate resident microglia via soluble factors, ultimately leading to demyelination^4^. Moreover, blood proteins have recently been shown to activate microglia also in the context of EAE^99^. Thus, upon their diffusion through the blood-brain barrier, soluble factors from plasma and/or CSF might be responsible for the widespread induction of glial disease-associated signatures. Lassmann and colleagues have previously proposed two distinct patterns of inflammation in MS, one leading to active cortical lesions and driven by soluble factors deriving from infiltrates in the meninges, and the other leading to slow expansion lesions and driven by soluble factors derived from perivascular space^10^. In accordance, central veins have been shown to associate with MRI-defined MS lesion^100, 101^. Since we observe that the global induction of DA glia is decoupled from active lesion formation, our results support the latter pattern.

The absence of homeostatic microglia during normal development has been recently shown to lead to acute hypermyelination and induction of a DA-oligodendrocyte state characterised by the expression of *Serpina3n* and *C4b*, followed by demyelination^102^. Thus, persistent conversion of homeostatic microglia into DA-microglia might *per se* be associated with the induction of a DA-oligodendrocyte state and ultimately also contribute to the demyelination observed in EAE. Given that DA-glia appears to be one of the earliest hallmarks of the disease, our data suggest that genes and proteins characterising these cell populations might be used as biomarkers of MS even prior to the development of symptoms.

Epstein-Barr virus infection of B cells and molecular mimicry have been gaining momentum as plausible triggers of MS^2, 103–105^. Interestingly, COVID-19 has also been suggested to exacerbate MS symptoms^106^, with the induction of DA-microglia in the CNS described as a hallmark of mild respiratory COVID-19 and H1N1 viral infections^107^. Since these viruses do not reach the CNS, increased levels of cytokines and chemokines in the CSF reportedly propelled the induction of DA microglia, which was global and persistent. In turn, this led to oligodendrocyte cell death and was ultimately associated with the brain “fog”, a common symptom of these diseases^107^. We hypothesise that the observed widespread DA-glial profile at peak EAE (**Figure 4**) is thus most likely triggered by the systemic release of cytokines from the vasculature. Consistent with these studies, we observed that microglia exhibited robust and long-lasting DA signatures along the disease course (**Figure 4**). Importantly, this was not observed in oligodendrocytes and astrocytes, which instead transitioned back to the homeostatic states outside late lesions (**Figure 4**), underlining the different glial responses to the same environment. Viral infections could thus act as initial players in MS aetiology mediating DA-glia states and oligodendrocyte cell death, which could then, combined with reactivation of EBV virus in B-cells and infiltration of adaptive immune cells in the CNS, lead to the development of disease. This option is not mutually exclusive with a possibility of molecular mimicry between Epstein Barr virus proteins and host CNS antigens such as Anoctamin 2^105^, GlialCam^103^ and Alfa B Crystallin^104^.

MS has been classically defined as relapsing-remitting, usually followed by primary progressive disease and secondary progressive, based on disease symptomatology and clinical screening of lesions in MS patients with Magnetic Resonance Imaging (MRI). MRI is one of the few technologies enabling spatiotemporal monitoring of MS lesions. Still, it has limited resolution to discriminate against different types of lesions^108^. Importantly, it is becoming increasingly clear that disease processes might affect normal-appearing tissue, which is not detectable by MRI^109^. This underlines the need for a deeper understanding of lesion formation and changes in transcriptional and cellular landscapes guiding these processes across anatomical regions, including normal appearing tissue along the course of the disease. The relapse-remitting and progressive disease definitions have recently been debated since progression does not entirely correlate with lesion load. Progression independent of relapse activity (PIRA) occurs already at early stages in the disease, also in relapsing-remitting patients^110^, and the smouldering course of the disease has been suggested to occur in parallel to lesion formation^109^. The occurrence of DA-glia within the EAE brain and the distribution of DA-glia throughout the parenchyma, also in lesion-deprived cervical regions **(Figure 4)**, is in line with the concept that MS progression is already present at the onset of the disease^109^.

During the last three decades, there has been remarkable progress in MS therapeutics, with anti-inflammatory agents and strategies targeting peripheral immune cells, such as B and T cells, alleviating the burden of the disease in patients with a relapsing-remitting course^111^. Nevertheless, success in treating progressive MS is limited, prompting the search for alternative therapeutic approaches. Investigation of the cellular and molecular mechanisms underlying MS has so far relied on the neuropathology and, recently, snRNA-Seq of human MS archival tissue and the modelling of specific disease aspects in mouse models. Our study, using ISS to investigate MS neuropathology in both an animal model of the disease and in human MS archival tissue at a spatial single-cell resolution, gives novel insights into the dynamics of MS disease evolution, uncoupling, for instance, global neuropathology from lesion evolution. By unveiling the interplay between different cell populations during the development of MS pathology and better understanding the cellular mechanisms underlying disease evolution within the CNS, our results uncover cellular and molecular pathways that can constitute novel targets for MS therapies.

### Limitations of the study

The development and application of technologies with single-cell and spatial resolution as ISS in this paper allows to investigate the disease in unprecedented resolution, which will lead to a better understanding of the disease mechanisms and aid in the development of novel therapeutic approaches. Nevertheless, MS is a complex and heterogeneous disease. The EAE model used in this study, while being the most prevalently investigated in the research community, is T-cell driven and while mimicking several aspects of MS pathology, it does not recapitulate aspects including the formation of brain lesions, B-cell contribution to the disease, presence of segregated inactive and remyelinating lesions, successive waves of inflammation among other aspects^112^. Thus, performing ISS in other mouse models that exhibit some of these other aspects of the disease will, in the future, better comprehend other aspects of the disease initiation and progression.

While providing unique spatial single-cell resolution, our study is limited to hundreds of marker genes that define different cellular populations. Even though we used markers for different subpopulations of glia, we were not able to discriminate between other glial cell states, indicating the need for additional markers. Using more probes while avoiding optical crowding, combining ISS with other orthogonal technologies such as DBit-Seq, MERFISH, CosMx/GeoMx, and Total-Seq might allow future investigation of cellular subpopulations at higher resolution and thus infer further aspects of diverse cellular contribution to MS.

## STAR MATERIALS

### KEY RESOURCES TABLE

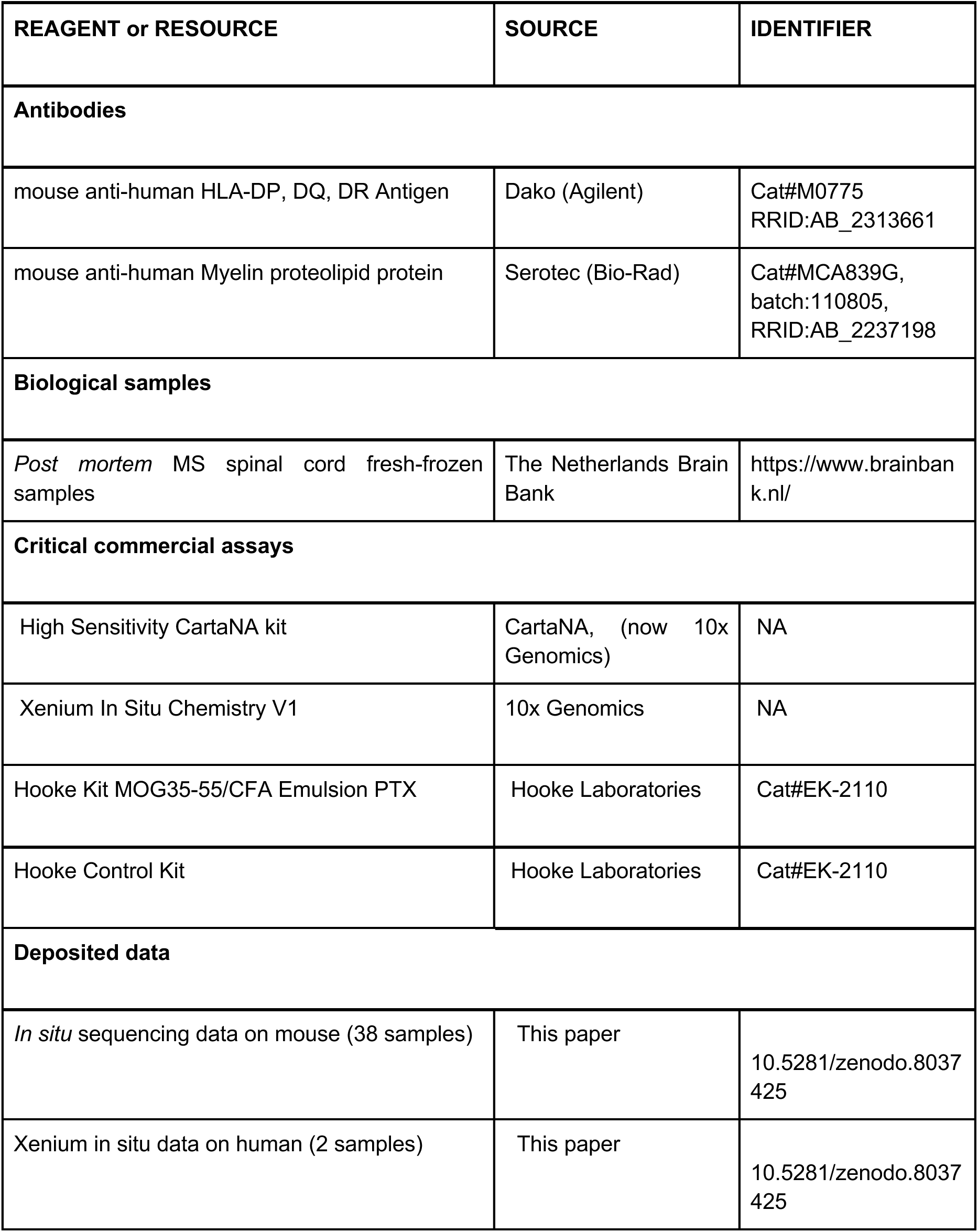
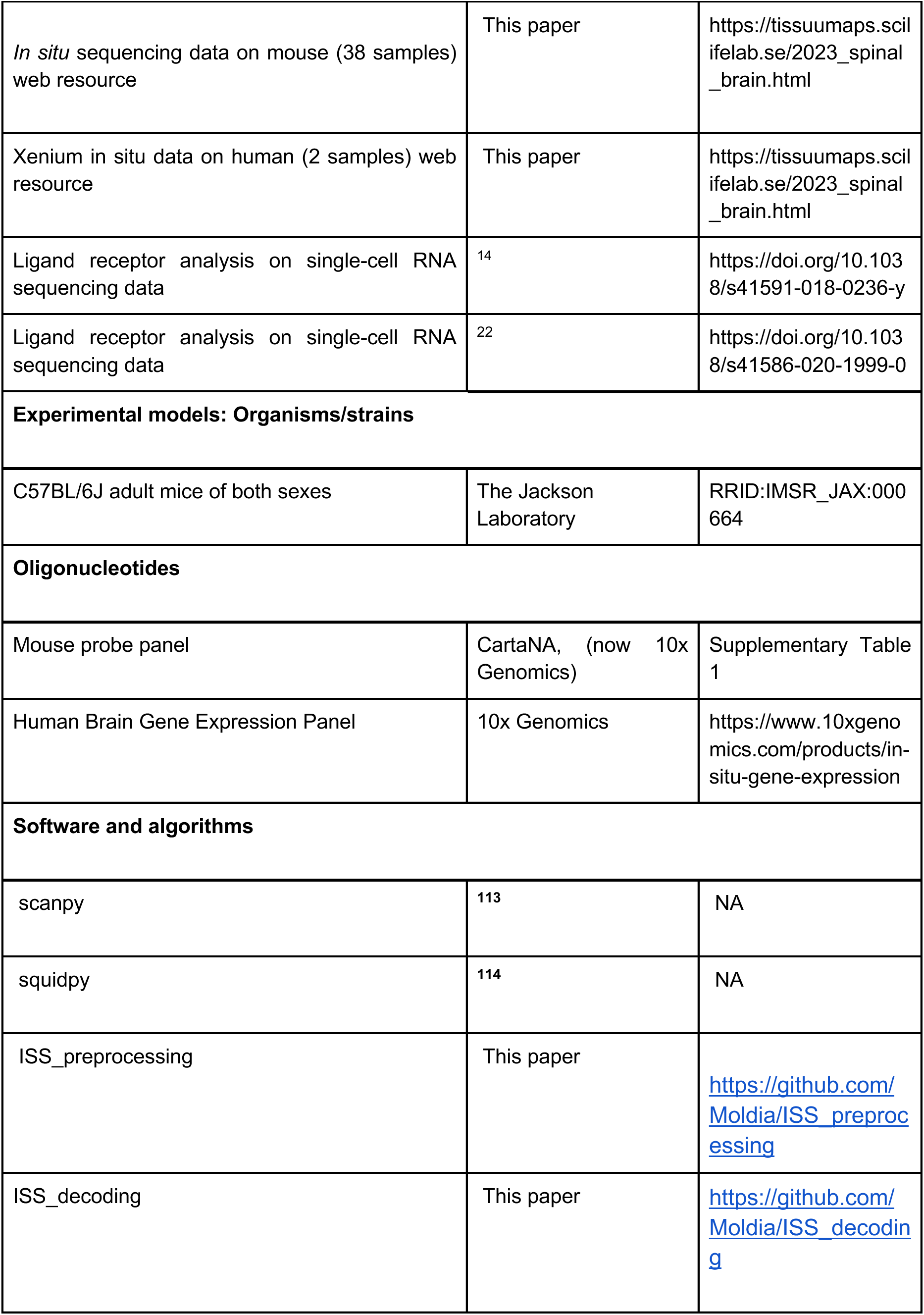
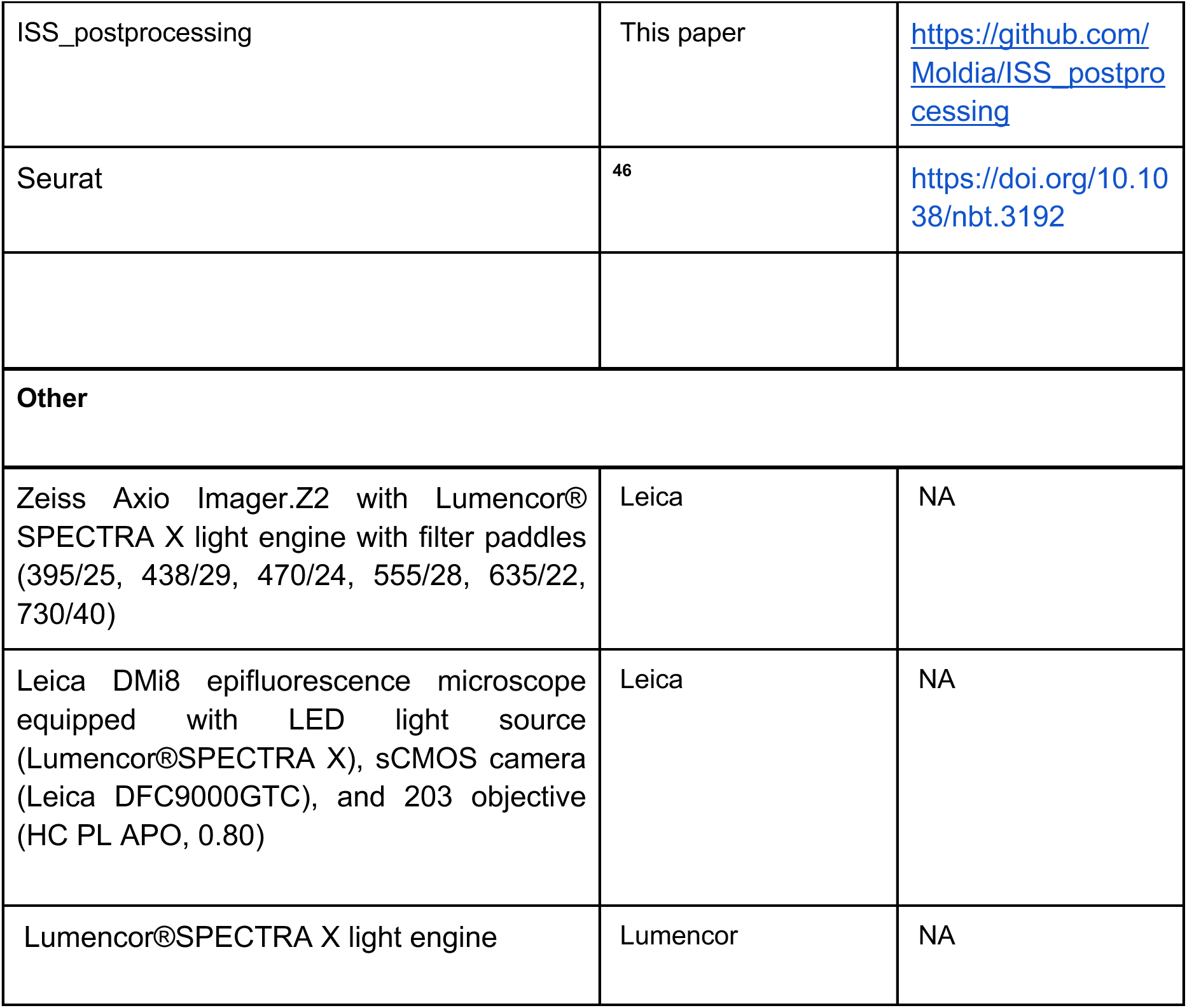

### RESOURCE AVAILABILITY

#### Lead contact

Further information and requests for resources and reagents should be directed to and will be fulfilled by the Lead Contact, Gonçalo Castelo-Branco.

#### Materials availability

The study did not generate new unique reagents.

#### Data code and availability

The ISS data from the 38 mouse samples and the two human MS samples are deposited at Zenodo under the DOI: 10.5281/zenodo.8037425. The data is available in one anndata format, compatible with scanpy and squidpy^115^. All of the code for the preprocessing of the microscopy images can be found at https://github.com/Moldia/ISS_preprocessing, decoding can be found at https://github.com/Moldia/ISS_decoding and the postprocessing at https://github.com/Moldia/ISS_postprocessing.

Spatial ISS datasets can be explored at: https://tissuumaps.scilifelab.se/2023_spinal_brain.html.

## EXPERIMENTAL MODEL AND SUBJECT DETAILS

### METHOD DETAIL

#### Animals and EAE experiments

C57BL/6J mice were purchased from the Jackson Laboratory and left to acclimate for at least one week before proceeding with the EAE experiments. All animals were free from mouse bacterial and viral pathogens, ectoparasites and endoparasites. The following light/dark cycle was maintained: dawn, 06:00–07:00; daylight, 07:00–18:00; dusk, 18:00–19:00; night, 19:00–06:00. Mice were housed in individually ventilated cages (IVC Sealsafe GM500, Tecniplast) at a maximum number of five per cage. General housing parameters, including temperature, ventilation and relative humidity, followed the *European Convention for the Protection of Vertebrate Animals used for experimental and other scientific purposes*. Air quality was controlled using stand-alone air-handling units equipped with a high-efficiency particulate air filter. Relative air humidity was consistently 55 ± 10%, at a temperature of 22 °C. Husbandry parameters were monitored with ScanClime (Scanbur) units. The cages contained a card box shelter, gnawing sticks and nesting material (Scanbur), placed on hardwood bedding (TAPVEI). Mice were provided with a regular chow diet, and hydrogel (Scanbur) was used as a supplement for the mice affected by EAE. Water was supplied, and water bottles were changed weekly. Cages were changed every second week in a laminar air-flow cabinet.

EAE was induced in male and female mice between the ages of 9 and 10 weeks. Mice from two independent EAE experiments were used in this study for peak and late EAE conditions. To induce EAE, mice were injected subcutaneously at two sites (proximal to cervical and lumbar spine) with an emulsion of MOG35-55 peptide in complete Freud’s adjuvant (CFA; MOG35-55/CFA Emulsion PTX (EK-2110), Hooke Laboratories), followed by intraperitoneal injection with pertussis toxin in PBS (200 ng per animal) on the day of immunisation and with a 24h delay (according to manufacturer’s instructions). Control animals underwent the same treatment, excluding the MOG35-55 peptide (Hooke Control Kit™ (CK-2110)). Spinal cord and brains were collected at the peak (one day after animals reached score 3, defined by the limp tail and complete paralysis of hind legs) and late stage of the disease (30 days after symptom onset, which was defined by the limp tail). Only animals with representative disease courses and clinical symptoms were used in this study.

Mice were sacrificed by anaesthesia with ketamine (120 mg kg–1 body weight) and xylazine (14 mg kg–1 body weight), followed by transcranial perfusion with cold oxygenated artificial cerebrospinal fluid (87 mM NaCl, 2.5 mM KCl, 1.25 mM NaH2PO4, 26 mM NaHCO3, 75 mM sucrose, 20 mM glucose, 1 mM CaCl2*2H2O and 2 mM MgSO4*7H2O in distilled H2O). Following isolation, brains and spinal cords were maintained for a minimal time period in the artificial cerebrospinal fluid until embedding in Tissue-Tek O.C.T. compound (Sakura) and snap-freezing using a mixture of dry ice and ethanol. Coronal cryosections of 10μm were mounted on Superfrost slides. The selection of SC tissue sections was specified by broad anatomical region but not the presence of lesions to ensure that representative sections were collected for ISS.

All experimental procedures were conducted following European directive no. 2010/63/EU, local Swedish directive no. L150/SJVFS/2019:9, Saknr no. L150 and Karolinska Institutet complementary guidelines for procurement and use of laboratory animals, no. Dnr. 1937/03-640. All procedures described were approved by the local committee for ethical experiments on laboratory animals in Sweden (Stockholms Norra Djurförsöksetiska nämnd, nos. 1995/2019 and 7029/2020).

#### Human tissue

The spinal cord *post mortem* archival samples were obtained from The Netherlands Brain Bank, Netherlands Institute for Neuroscience, Amsterdam (open access: www.brainbank.nl). Two fresh-frozen cervical mirror blocks (https://www.brainbank.nl/nbb-ms/characterization-of-ms-lesions/) of the ones with pathologically annotated lesions were used for ISS analysis. The sample with active lesions from a female donor, age 53, with a *post mortem* collection interval of 7h15min and stored at -80 for 10 years. The sample with inactive lesions from a male donor, age 47, with a *post mortem* collection interval of 4h25min and was stored at -80 for 16 years. All Material has been collected from donors for or from whom written informed consent for a brain autopsy, and the use of the material and clinical information for research purposes had been obtained by the NBB. Human archival tissue was used in accordance with the ethical permit approved by the Swedish Ethical Review Authority (Stockholm, Sweden, reference no. 2016/589-31 with amendment 2019-01503).

#### Mouse panel probe selection

Genes were selected to delineate cell types in the EAE model. That included main populations of immune cells, neurons, astrocytes, oligodendrocytes, microglia, vascular cells and, within each population, a subset of cell types. This selection was made manually in combination with different computational frameworks^116, 117^. The gene list was sent to CartaNA AB (part of 10x Genomics), which designed the padlock probes. The target sequences and design of the padlock probes are proprietary information.

#### *In situ* sequencing assay

The reagents and protocol were supplied by CartaNA AB. First, the tissue samples were fixed and after fixation, the dRNA probe mixture was added and left to incubate overnight at 37°C. The following day, the ligation mixture was added and left to incubate at 37°C for 2 hours. Following the ligation, the rolling circle amplification was performed overnight at 30 °C. The rolling circle products were then read out in situ by microscopic imaging.

#### Xenium In Situ Gene Expression

Fresh frozen 10μm tissue sections were placed on Xenium slides. The tissue was fixed and permeabilized as described in Xenium Fixation and Permeabilization Protocol (Demonstrated protocol CG000581). Predesign human brain panel was added to the tissue (266 genes, chemistry version 1: https://cdn.10xgenomics.com/raw/upload/v1684952565/software-support/Xenium-panels/hBrain_panel_files/Xenium_hBrain_v1_metadata.csv). Probes were hybridised to the target RNA, ligated, and enzymatically amplified generating multiple copies for each RNA target as described in Probe Hybridization, Ligation and Amplification user guide (User guide CG000582). Xenium slides were then loaded for imaging and analysis on the Xenium Analyzer instrument following the Decoding and Imaging user guide (User guide CG000584). Instrument software version 1.3.3.0 and software analysis version 1.3.0.6 were used.

#### Imaging

The imaging was performed using an epifluorescence microscope of make and model Zeiss Axio Imager.Z2 and LEICA DMi8. The two microscopes were connected to external LED sources, Lumencor® SPECTRA X light engine and Lumencor LED light engine SpectraX, respectively. The two systems were equipped with filter paddles (395/25, 438/29, 470/24, 555/28, 635/22, 730/40), and images were taken with sCMOS cameras (2048 × 2048, 16-bit, ORCA-Flash4.0 LT Plus, Hamamatsu and Leica DFC90000GTC-VSC10726). To separate wavelengths, filter cubes were used. Specifically, quad band Chroma 89402 (DAPI, Cy3, Cy5), quad band Chroma 89403 (Atto425, TexasRed, AlexaFluor750), and single band Zeiss 38HE (AlexaFluor488) as well as AlexaFluor 488/FITC/FAM dyes; SP102v2 (Chroma) for imaging Cy3; SP103v2 (Chroma) for imaging Cy3.5/TexasRed; SP104v2 (Chroma) for imaging Cy5; 49007 (Chroma) for imaging Cy7/AlexaFluor 750 dyes on the LEICA system. An automatic multi-slide stage was used (automatic multi-slide stage (LMT200-HS). The imaging was done in tiles (2048×2048 pixels) with a 10% overlap and 24 z-stack planes with 0.5μm spacing. Raw tiff files with accompanying metadata files were exported.

#### Image deconvolution and restoration

To increase the number of decodable spots in dense imaging data, we first used Deconwolf^39^ on 3 sections in order to generate a training set for image restoration using CARE^40^.

## QUANTIFICATION AND STATISTICAL ANALYSIS

### Image processing and decoding of HybISS data

#### Preprocessing

The preprocessing of the images from the microscope included maximum intensity projection of the z stacks, alignment of tiles between cycles and image stitching. The code can be found at https://github.com/Moldia/ISS_preprocessing. In short, the images and the accompanying metadata were exported during the imaging. Images were projected and formatted into OME tif files and subsequently stitched and aligned using ASHLAR ^118^. Lastly, the stitched images were subsequently sliced 2000 by 2000 pixels to reduce the computational requirements for decoding.

#### Decoding

The decoding of transcripts is based on the python package called starfish (https://spacetx-starfish.readthedocs.io/en/latest/). The code can be found at https://github.com/Moldia/ISS_decoding. In short, images are registered and white top hat filtered. Then the channel intensities are normalised across channels and spots are located in a composite maximum intensity projected image of signal images from the same field-of-view and sequencing round. The spots were decoded using PerRoundMaxChannel, in which, for each spot and each base, the highest intensity channel was determined and matched to a corresponding barcode in the codebook. In addition, a quality metric was assigned to every spot in every cycle, defined as the called channel intensity divided by the sum of all other channels (between 0.25 and 1).

#### Cell segmentation and assigning spots to cells

Cells were segmented using StarDist^41^. We manually drew and annotated cells to generate a training set of images to refine the base model (2D_versatile_fluo). This was done since we noted missegmentations in the dense lesion areas. This was done on resliced images, and the segmentation mask for each field-of-view was subsequently stitched together, and each cell was labelled with a unique identifier. Metadata about the cells were extracted (x and y coordinate and area). Transcripts were then assigned to cells. Annotated data objects were created^119^.

#### De novo clustering

We proceeded by retaining cells with at least 5 transcripts and 3 genes expressed. Following filtering, the gene expression was normalised, transformed and scaled, and the gene expression content of each cell was then clustered using Leiden clustering^120^. In this process, we worked in a hierarchical manner by first doing a coarse clustering to identify the broad populations of macro- and microglia, neurons, immune cells and vascular cells. Once these broad clusters had been annotated, the broad clusters were subclustered to define fine clusters.

#### Xenium processing

The human *post mortem* MS spinal cord data was generated on the newly released Xenium In Situ platform by the In Situ Sequencing facility at Science for Life Laboratory in Solna, Stockholm. The panel used in the assay was the newly released Human Brain Panel from 10x genomics, probing for 266 genes. We generated gene expression matrices from the Xenium output by considering only transcripts closer than 5 µm away from the nucleus. These expression matrices were subsequently processed in the scanpy framework. We considered cells with more than 10 transcripts and 3 genes expressed. We employed the same strategy for cell type identification as outlined above. In short, the gene expression was normalised, transformed, scaled and subsequently clustered using Leiden clustering. We used a hierarchical strategy here as well, firstly identifying broad clusters that were subsequently subclustered.

#### Annotation of compartments

The compartments in the mouse and human data were defined computationally by looking at the local neighbourhood (maximum distance of 100 µm) around each cell. This generated a neighbourhood matrix that was subsequently clustered and annotated.

#### Gene module scores

When computing the gene module score we used the sc.tl.score_genes function in scanpy. The scoring process involves comparing the average expression level of a specific set of genes with the average expression level of a reference set of genes. The reference set is generated by randomly selecting genes from the larger pool of available genes, taking into account the binned expression values. By subtracting the average expression of the reference set from the average expression of the target gene set, a score is obtained for each cell. For the IFN-gamma response we used *Irf7, Irgm1, Irgm2, Igtp, Zbp1*, and the Serpins gene modules comprised of *Serpina3n, Serpina3h, Serpina3c, Serpina3i, Serpina3f, Serping1, Serpina3m*. For the major histocompatibility complex class of genes, we used for class 1: *H2-K1, H2-D1, H2-T23, B2m, Psmb9, Tap1, Nlrc5* and for class 2: *Cd74, H2-Aa, H2-Ab1, H2-Eb1, H2-DMa, Ctss, Ciita*. For the myelination module, we used *Apod, 9630013A20Rik, Egr2, Pmp22*.

#### Neighbourhood analysis in squidpy

For the neighbourhood analysis in Squidpy, we first computed the spatial neighbourhood graph, connecting the 5 nearest components for each cell. Then we computed two metrics, the neighbourhood enrichment score and the spatial connectivities. These two metrics are complementary in the sense that the spatial connectivities calculate the cumulative count of connections between different cell types. The enrichment score is determined by analysing the proximity of cell clusters on a connectivity graph. It compares the observed events to a randomly permuted dataset and calculates a z-score to assess the statistical significance.

### Inference of cell-cell communications from scRNA-seq

Inference of receptor-ligand communication was performed using the published single RNA-seq expression matrix and annotations: GSE130119^22^, GSE113973^14^ (expression matrix available under request by the authors). Only CFA and EAE peak cells were selected from both studies. Both matrices were processed and integrated in R 4.1.3 using Seurat 4.3.0^121^, with parameters 2000 variable features, *SelectIntegrationFeatures and FindIntegrationAnchors*. The clusters were annotated according to the ISS defined clusters. First, a gene module score using the top differentially expressed genes of the ISS annotated celltypes was calculated, with parameters *AddModuleScore ctrl=15* from Seurat. Module scores were binarized based on their distribution. By combining binary module scores, the original annotations from the published studies and canonical markers the cells were assigned to ISS celltype annotations. Most of the cells corresponded to their original published broad celltype classification. Clusters with few cells were removed for receptor-ligand inference.

Selected scRNA-seq ISS clusters were downsampled to a maximum of 50 cells per cluster, to avoid population size biases. Due to the small number of cells, Neurons and NFOL clusters were removed. Downsampled Seurat object was used as input for CellChat 1.6.1 R package **(REF)** with parameters *computeCommunProb(type= “triMean”, population.size = F* and *filterCommunication(min.cells = 10)*. Visualization of the significant communications (p-val < 0.01) for DA_MOL56 and DA_MOL2 was done using the function *netVisual_bubble from CellChat*.

## Acknowledgements

We thank Tony Jimenez-Beristain for writing the laboratory animal ethical permits 1995_2019 and 7029/2020 and for the assistance with animal experiments. We thank Chao Zheng, Karl Carlström and Celine Geywitz for advice and help with experimentation, and Yonglong Dang for advice on human MS sample selection. We thank Ting Sun for critically evaluating the manuscript. We thank Rasmus Berglund for the help with the microglia marker panel selection and Sergio Marco Salas and Marco Grillo for valuable input on the neighbourhood analysis and the image restoration. We thank The Netherlands Brain Bank (in particular Isabell Ehmer and Michiel Kooreman), the donors and their families for the MS archival tissue used in this tissue. We acknowledge support from the *In situ* sequencing unit in Stockholm funded by Science for Life Laboratory, the National Genomics Infrastructure in Stockholm funded by Science for Life Laboratory, the Knut and Alice Wallenberg Foundation and the Swedish Research Council. Part of the computation/data handling was enabled by resources provided by the National Academic Infrastructure for Supercomputing in Sweden (NAISS) and Swedish National Infrastructure for Computing (SNIC) at the Uppsala Multidisciplinary Center for Advanced Computational Science, partially funded by the Swedish Research Council through grant agreement no. 2022-06725 and no. 2018-05973. Part of the computing was also performed in the Linnarsson group Monod Linux cluster at MBB-KI, and we thank Peter Lönnerberg for maintenance and support. Work in M.N.’s research group was supported by Chan Zuckerberg Initiative, an advised fund of Silicon Valley Community Foundation (DAF2018-191929); Erling-Persson Family Foundation (A human developmental cell atlas); Knut and Alice Wallenberg Foundation (KAW 2018.0172); Swedish Research Council (2019-01238) and Swedish Cancer Society (Cancerfonden; CAN 2021/1726). M.N. and G.C.-B. were co-funded by a grant from the Strategic Research Programme in Neuroscience (StratNeuro). Work in G.C.-B.’s research group was supported by the Swedish Research Council (grant 2019-01360), the European Union (Horizon 2020 Research and Innovation Programme/ European Research Council Consolidator Grant EPIScOPE, grant agreement number 681893), the Swedish Brain Foundation (FO2018-0162), the Swedish Cancer Society (Cancerfonden; 190394 Pj), Knut and Alice Wallenberg Foundation (grants 2019-0107 and 2019-0089), the Göran Gustafsson Foundation for Research in Natural Sciences and Medicine, the Swedish Society for Medical Research (SSMF, grant JUB2019), Olav Thon Foundation, Ming Wai Lau Centre for Reparative Medicine, and Karolinska Institutet.

## Author contributions

Conceptualization, P.K., C.M.L., M.N. & G.C.-B.; Methodology, C.M.L., P.K, L.A.R.R.-K., M.M.H., M.N. & G.C.-B.; Software, C.M.L., E.A., C.A.; Formal Analysis, C.M.L., P.K., E.A.; Investigation, P.K., A.R., C.Y., K.T.; Resources, P.K., C.M.L., L.A.R.R.-K, E.A., A.R., C.Y, C.A., K.T., A.O.G.-C., M.M.H.; Data Curation, C.M.L; Visualization, C.M.L., P.K., C.A.; Supervision, L.A.R.R.-K., A.O.G.-C., T.O., M.M.H., M.N. & G.C.-B.; Writing – Original Draft, P.K., G.C.-B., C.M.L., M.M.H. & M.N.; Funding Acquisition, T.O., M.N. & G.C.-B.. C.M.L. and K.P. have equal contributions in this paper, which should be cited with BOTH their surnames. The order of their names is interchangeable, and the paper can therefore be cited either as Langseth & Kukanja et al. or Kukanja & Langseth et al..

