## Supplementary Figures for "Single cell-resolution *in situ* sequencing elucidates spatial dynamics of multiple sclerosis lesion and disease evolution"

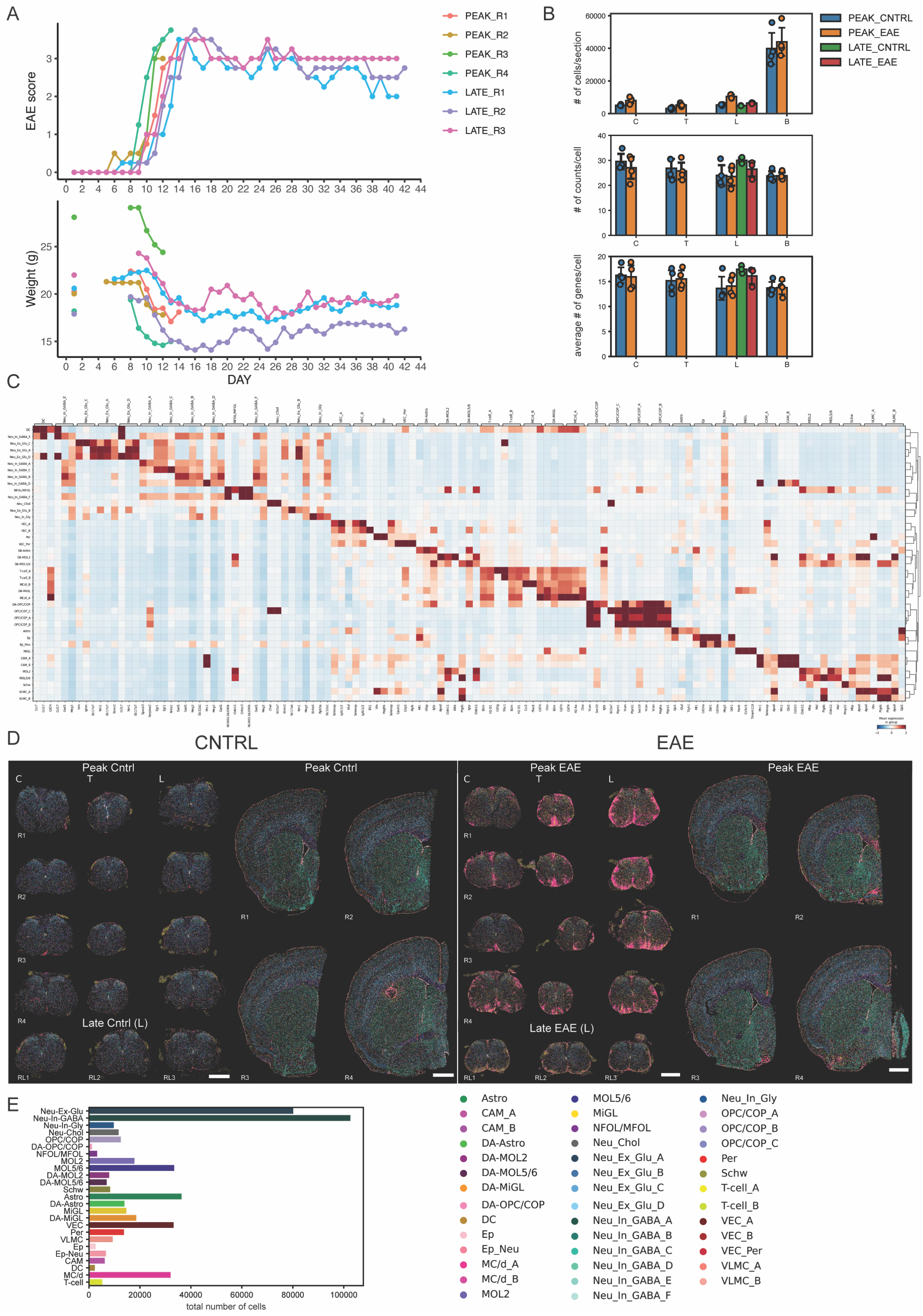

A
EAE score
Weight (g)
DAY
PEAK_R1
PEAK_R2
PEAK_R3
PEAK_R4
LATE_R1
LATE_R2
LATE_R3
B
### of cells/section
### of counts/cell
average # of genes/cell
PEAK_CNTRL
PEAK_EAE
LATE_CNTRL
LATE_EAE
C
D
CNTRL
EAE
Peak Cntrl
Late Cntrl (L)
Peak EAE
Late EAE (L)
E
total number of cells
Neu-Ex-Glu
Neu-In-GABA
Neu-In-Gly
Neu-Chol
OPC/COP
DA-OPC/COP
NFOL/MFOL
MOL2
MOL5/6
DA-MOL2
DA-MOL5/6
Schw
Astro
DA-Astro
MiGL
DA-MiGL
VEC
Per
VLMC
Ep
Ep-Neu
CAM
DC
MC/d
T-cell
Astro
CAM_A
CAM_B
DA-Astro
DA-MOL2
DA-MOL5/6
DA-MiGL
DA-OPC/COP
DC
Ep
Ep_Neu
MC/d_A
MC/d_B
MOL2
MOL5/6
MiGL
NFOL/MFOL
Neu_Chol
Neu_Ex_Glu_A
Neu_Ex_Glu_B
Neu_Ex_Glu_C
Neu_Ex_Glu_D
Neu_In_GABA_A
Neu_In_GABA_B
Neu_In_GABA_C
Neu_In_GABA_D
Neu_In_GABA_E
Neu_In_GABA_F
Neu_In_Gly
OPC/COP_A
OPC/COP_B
OPC/COP_C
Per
Schw
T-cell_A
T-cell_B
VEC_A
VEC_B
VEC_Per
VLMC_A
VLMC_B

***Supplementary Figure 1, QC metrics and replicates of ISS EAE experiments, related to Figure 1.***

***(A)*** *Disease scores and corresponding body weights of EAE mice whose CNS were used in ISS.*

***(B)*** *The mean numbers of cells per section, read counts per cell, and genes per cell across different experimental conditions. C=cervical spinal cord, T=thoracic spinal cord, L=lumbar spinal cord, B=brain (hemisphere).*

***(C)*** *Heatmap of top DE genes for high-resolution clustering.*

***(D)*** *Spatial representation of the 41 annotated clusters (high-resolution clustering) in CNTRL and EAE spinal cord and brain samples. Scale bars - 1000µm. C=cervical spinal cord, T=thoracic spinal cord, L=lumbar spinal cord, R=replicate (corresponding to (A)). For exploration: https://tissuumaps.scilifelab.se/2023_spinal_brain.html.*

***(E)*** *Broad clusters with corresponding total numbers of cells (from control and EAE conditions combined).*

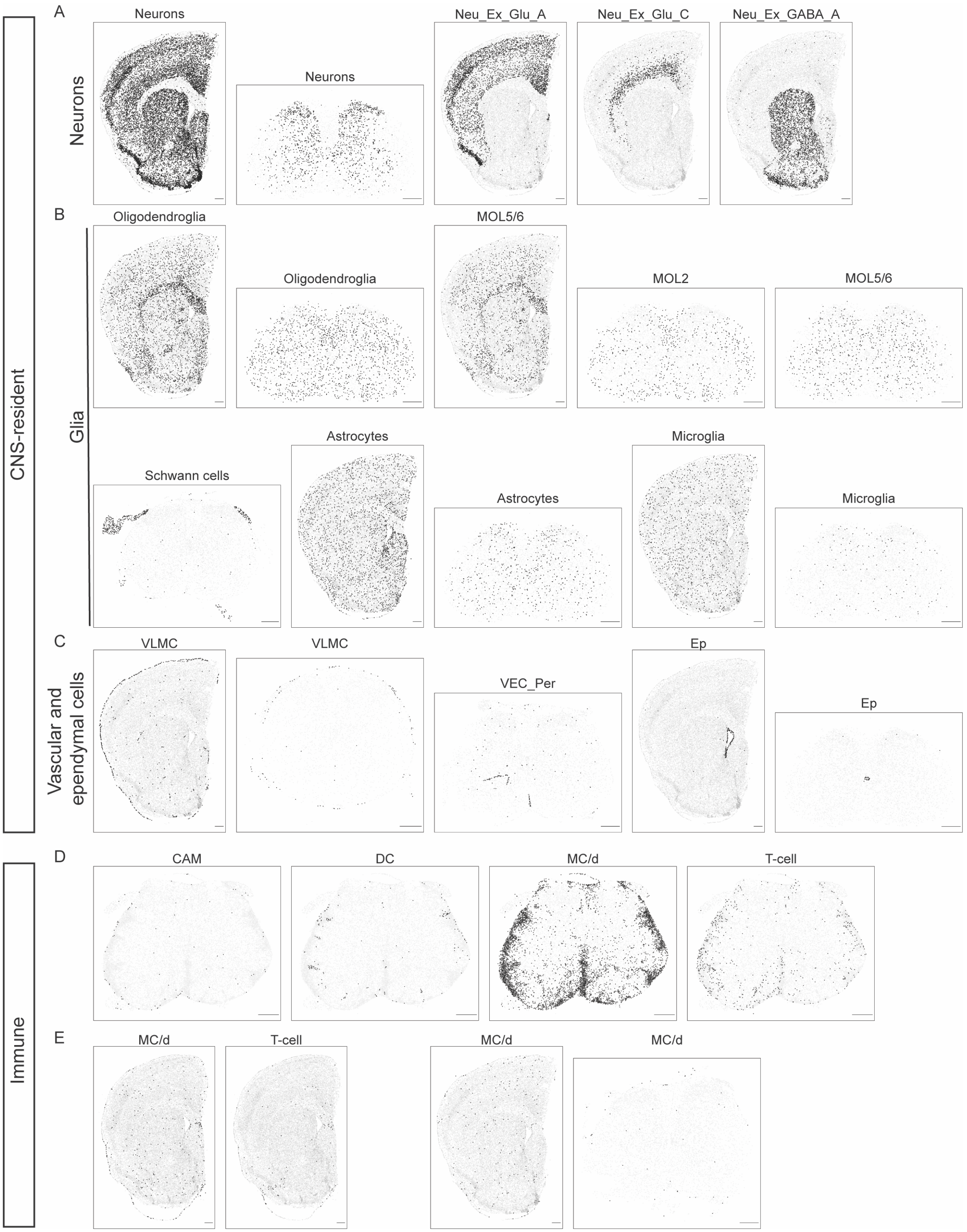
A
B
C
D
E
CNS-resident
Immune
Neurons
Glia
Vascular and ependymal cells
Neurons
Neurons
Neu_Ex_Glu_A
Neu_Ex_Glu_C
Neu_Ex_GABA_A
Oligodendroglia
Oligodendroglia
MOL5/6
MOL2
MOL5/6
Schwann cells
Astrocytes
Astrocytes
Microglia
Microglia
VLMC
VLMC
VEC_Per
Ep
Ep
CAM
DC
MC/d
T-cell
MC/d
T-cell
MC/d
MC/d

***Supplementary Figure 2, Spatial maps of homeostatic neural and immune populations, related to Figure 1.***

***(A-C)*** *Representative spatial maps from controls depicting selected homeostatic populations of neurons, glia, vascular and ependymal cells. Scale bars - 325μm. Neurons=total neuronal population, Ex=excitatory, Glu=glutamatergic, GABA=gabaergic, Oligodendroglia=total oligodendroglial lineage cells, MOL= mature oligodendrocytes, Astrocytes=total astrocyte population, Microglia=total microglial population, VLMC=vascular leptomeningeal cells, VEC=vascular endothelial cells, Per=pericytes, Ep=ependymal cells.*

***(D-E)*** *Representative spatial maps from peak EAE depicting selected immune populations. Scale bars - 325μm. CAM=CNS-associated macrophages, DC=dendritic cells, MC/d = monocytes and monocyte-derived cells.*

A

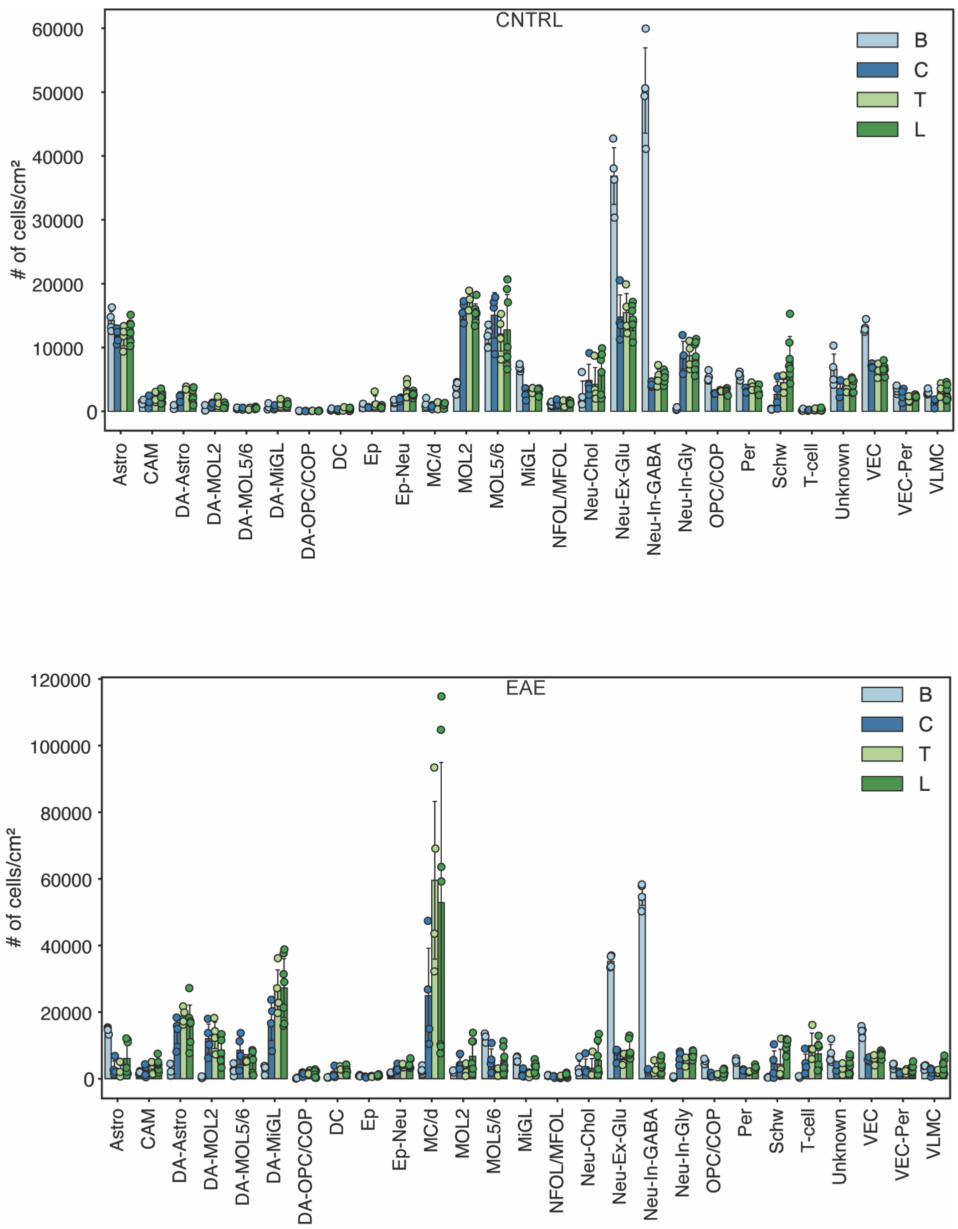

CNTRL
EAE
B
C
T
L
### of cells/cm²
Astro
CAM
DA-Astro
DA-MOL2
DA-MOL5/6
DA-MiGL
DA-OPC/COP
DC
Ep
Ep-Neu
MC/d
MOL2
MOL5/6
MiGL
NFOL/MFOL
Neu-Chol
Neu-Ex-Glu
Neu-In-GABA
Neu-In-Gly
OPC/COP
Per
Schw
T-cell
Unknown
VEC
VEC-Per
VLMC


B

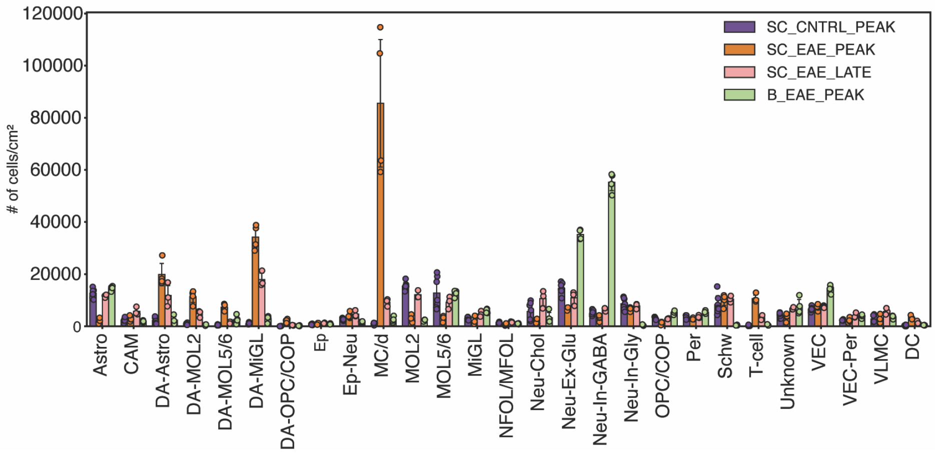

SC_CNTRL_PEAK
SC_EAE_PEAK
SC_EAE_LATE
B_EAE_PEAK
### of cells/cm²
Astro
CAM
DA-Astro
DA-MOL2
DA-MOL5/6
DA-MiGL
DA-OPC/COP
Ep
Ep-Neu
MC/d
MOL2
MOL5/6
MiGL
NFOL/MFOL
Neu-Chol
Neu-Ex-Glu
Neu-In-GABA
Neu-In-Gly
OPC/COP
Per
Schw
T-cell
Unknown
VEC
VEC-Per
VLMC
DC

***Supplementary Figure 3, Quantification of number of cells per cm² across regions and time points in EAE, related to Figure 1.***

***(A)*** *Number of cells per cm² for the identified cell populations in controls and EAE along the four different tissue types, brain (B), cervical (C), thoracic (T) and lumbar (L) spinal cord.*

***(B)*** *Number of cells per cm² for the identified cell populations in controls and EAE along the two different time points.*

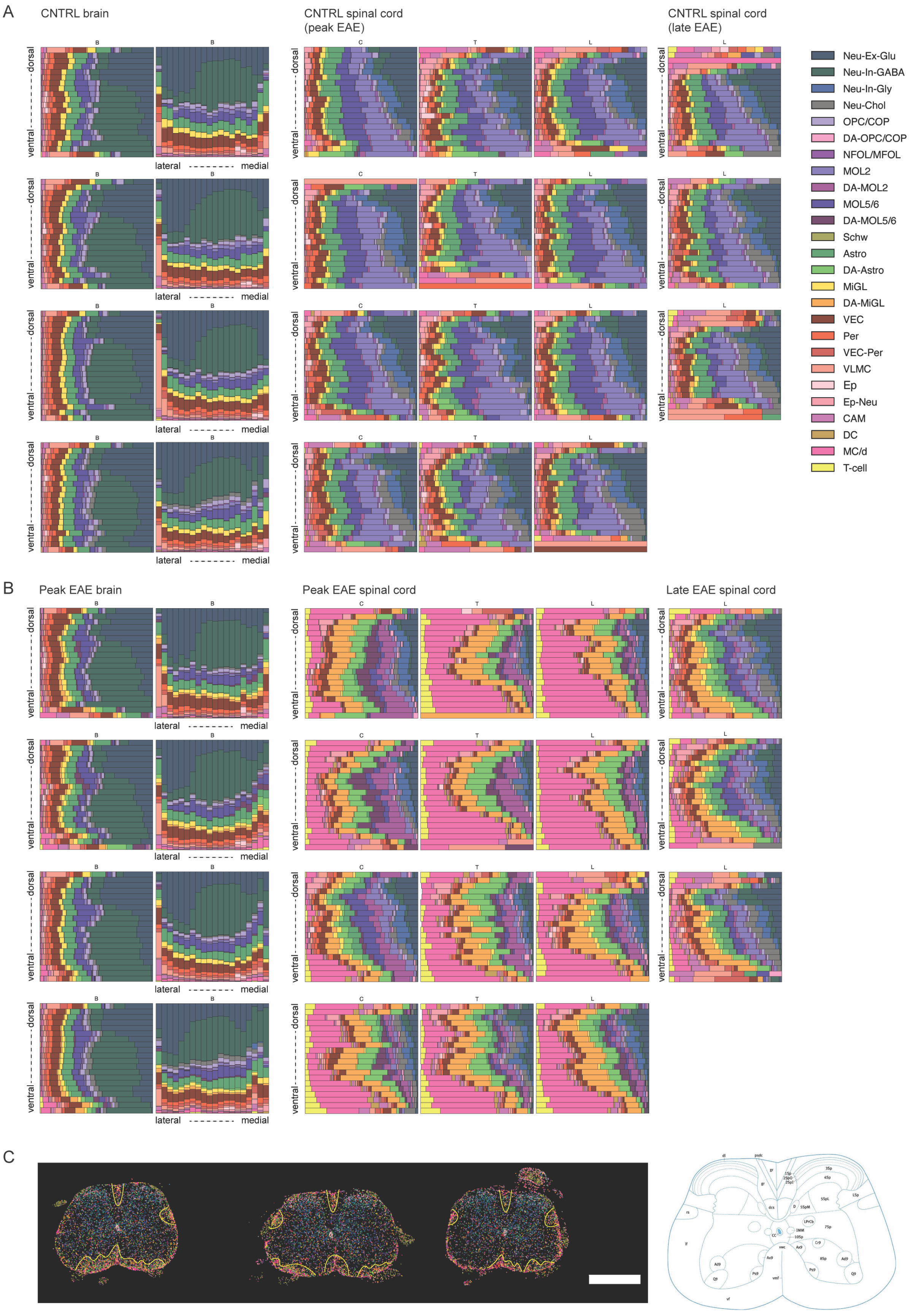
A
CNTRL brain
CNTRL spinal cord (peak EAE)
CNTRL spinal cord (late EAE)
B
C
T
L
dorsal
ventral
lateral
medial
Neu-Ex-Glu
Neu-In-GABA
Neu-In-Gly
Neu-Chol
OPC/COP
DA-OPC/COP
NFOL/MFOL
MOL2
DA-MOL2
MOL5/6
DA-MOL5/6
Schw
Astro
DA-Astro
MiGL
DA-MiGL
VEC
Per
VEC-Per
VLMC
Ep
Ep-Neu
CAM
DC
MC/d
T-cell
B
Peak EAE brain
Peak EAE spinal cord
Late EAE spinal cord
C

***Supplementary Figure 4, Dorsal-to-ventral and lateral-to-medial cell gradients in EAE, related to Figure 1.***

***(A)*** *The dorsal-to-ventral (brain and spinal cord) and lateral-to-medial (brain) gradients of cell types in controls. C=cervical spinal cord, T=thoracic spinal cord, L=lumbar spinal cord, B=brain (hemisphere).*

***(B)*** *The dorsal-to-ventral (brain and spinal cord) and lateral-to-medial (brain) gradients of cell types in EAE samples from the peak and late stage. C=cervical spinal cord, T=thoracic spinal cord, L=lumbar spinal cord, B=brain (hemisphere). See colour legend in A.*

***(C)*** *Spatial cell maps of lumbar spinal cord sections from late EAE with highlighted immune-infiltrated regions that correspond to distinct spinal cord anatomical regions depicted on the right (source: Allen Brain Atlas, https://mouse.brain-map.org). See colour legend in A. Scale bar - 1000µm.*

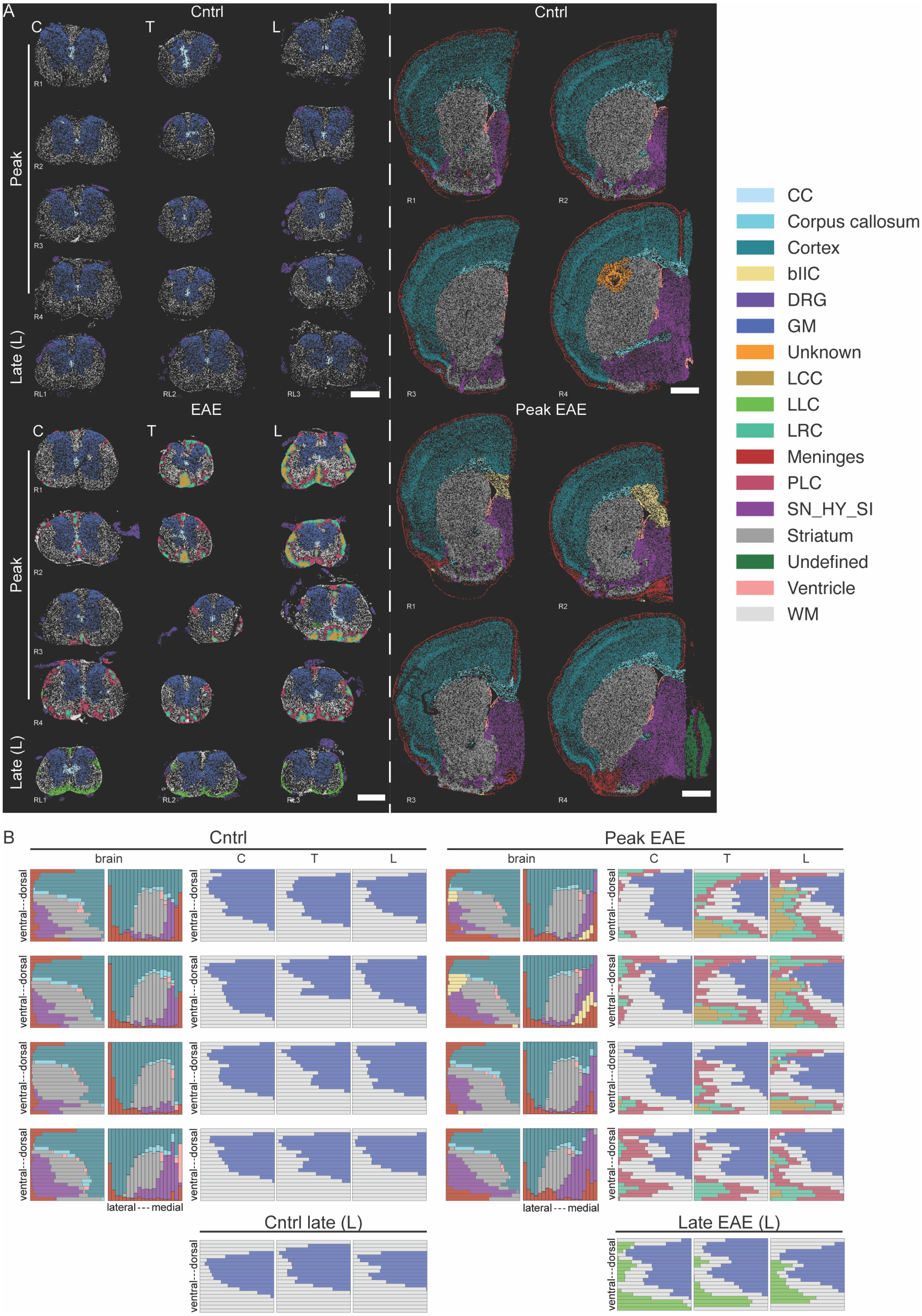

A
Cntrl
C
T
L
Peak
Late (L)
R1
R2
R3
R4
RL1
RL2
RL3
EAE
Peak EAE
CC
Corpus callosum
Cortex
bIIC
DRG
GM
Unknown
LCC
LLC
LRC
Meninges
PLC
SN_HY_SI
Striatum
Undefined
Ventricle
WM
B
brain
ventral---dorsal
lateral --- medial
Cntrl late (L)
Late EAE (L)

***Supplementary Figure 5, Spatial representation and dorsal-to-ventral and lateral-to-medial gradients of EAE compartments, related to Figure 2.***

***(A)*** *Spatial representation of the automated compartment annotations. Scale bars - 1000μm. R refers to "biological replicate", C refers to cervical, T to thoracic and L to lumbar spinal cord. Brain hemispheres are depicted on the right. For exploration: https://tissuumaps.scilifelab.se/2023_spinal_brain.html.*

***(B)*** *Comparison of compartment occurrences in control, peak and late EAE sections across dorsal-to-ventral (brain hemispheres and spinal cords) and lateral-to-medial (brain hemispheres) axes. The mean number of cells for each given compartment, normalised by the area of the section is shown. Number of bins = 20 in each section.*

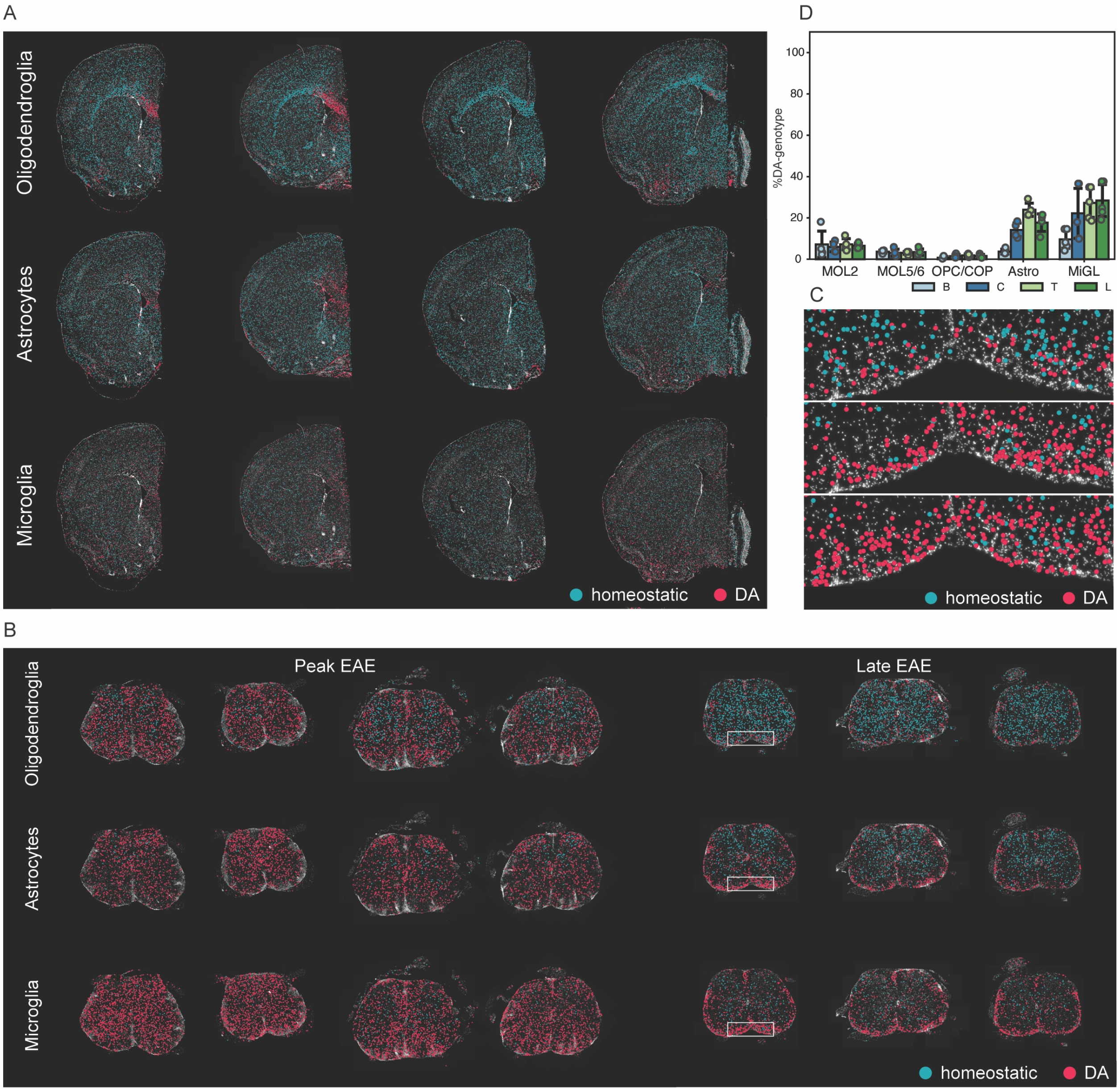


***Supplementary Figure 6, Spatial distribution of homeostatic and disease-associated glia, related to Figure 4.***

***(A)*** *Spatial distribution of homeostatic and disease-associated oligodendroglia, astrocytes and microglia across the four replicates of EAE brain.*

***(B)*** *Spatial distribution of homeostatic and disease-associated oligodendroglia, astrocytes and microglia across the four lumbar spinal cord replicates from peak EAE and three lumbar spinal cord replicates from late EAE.*

***(C)*** *Insets from* ***B*** *showing zoom-ins of the late lesion compartment and the distribution of homeostatic and disease-associated oligodendroglia (top), astrocytes (middle) and microglia (bottom).*

*(**D)** The percentages of disease-associated MOL2, MOL5/6, astrocytes (Astro) and microglia (MiGL) in control brain (B), cervical (C), thoracic (T) and lumbar (L) spinal cord sections.*

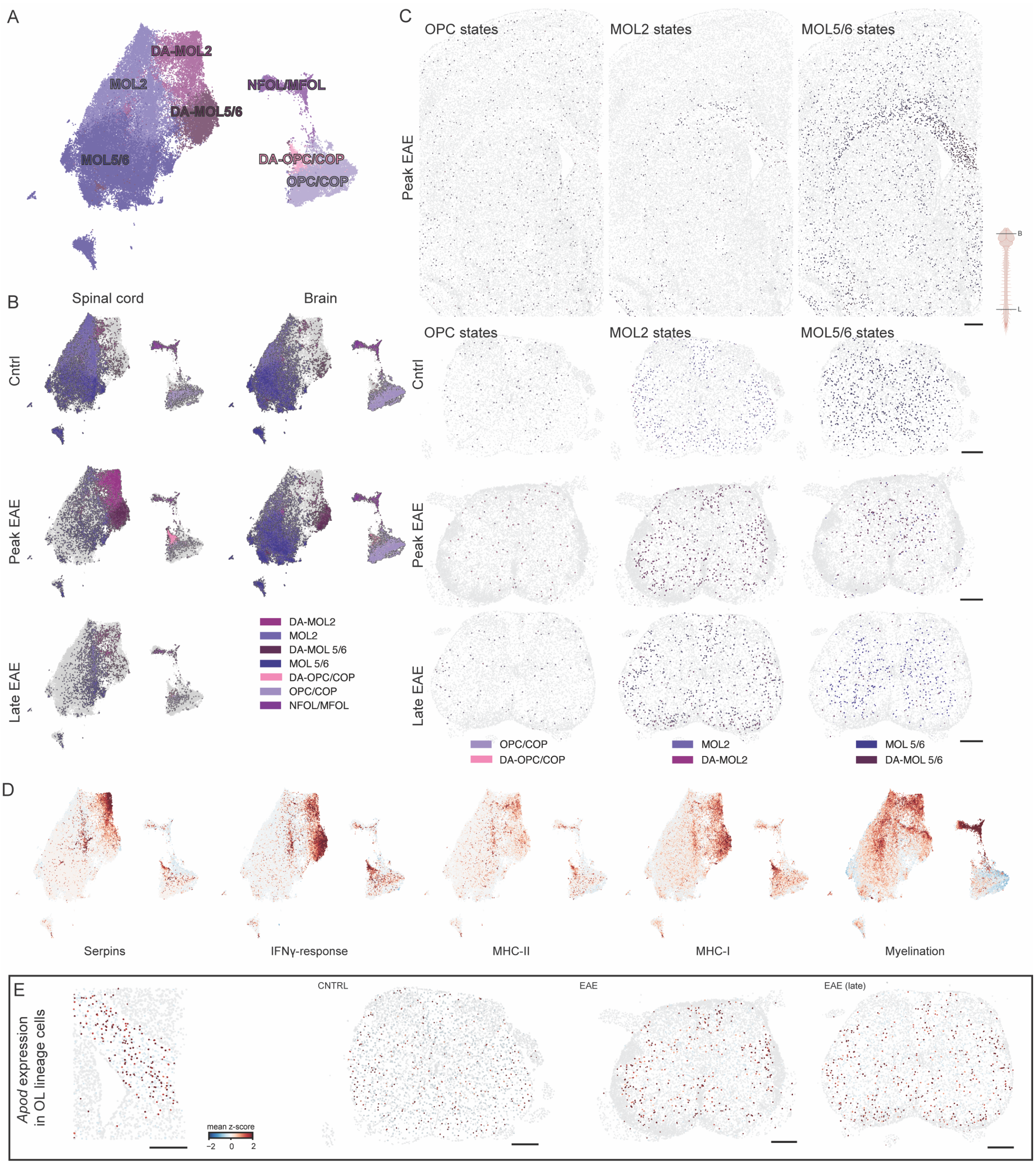


***Supplementary Figure 7, Spatial distribution of homeostatic and disease-associated oligodendroglia, related to Figure 4.***

***(A)*** *UMAP depicting subclusters of oligodendrocyte lineage cells.*

***(B)*** *UMAP representation of cells corresponding to different conditions, spinal cord in control,* *peak and late EAE and for brain, control and peak EAE.*

***(C)*** *Spatial maps of homeostatic and disease-associated oligodendrocyte lineage cells in representative sections of EAE brain (peak), control, peak EAE and late EAE lumbar spinal cord. Scale bars - 325μm.*

***(D)*** *UMAP depicting gene expression modules in OL lineage cells. For the "IFNg-response" module Irf7, Irgm1, Irgm2, Igtp, Zbp1 were used, and the "Serpins" gene module comprised of Serpina3n, Serpina3h, Serpina3c, Serpina3i, Serpina3f, Serping1, Serpina3m. The major histocompatibility complex modules of included for class 1: H2-K1, H2-D1, H2-T23, B2m, Psmb9, Tap1, Nlrc5 and for class 2: Cd74, H2-Aa, H2-Ab1, H2-Eb1, H2-DMa, Ctss, Ciita. For the myelination module, Apod, 9630013A20Rik, Egr2, and Pmp22 were used.*

***(E)*** *Expression of Apod in oligodendrocyte lineage cells. Scale bars - 325μm.*

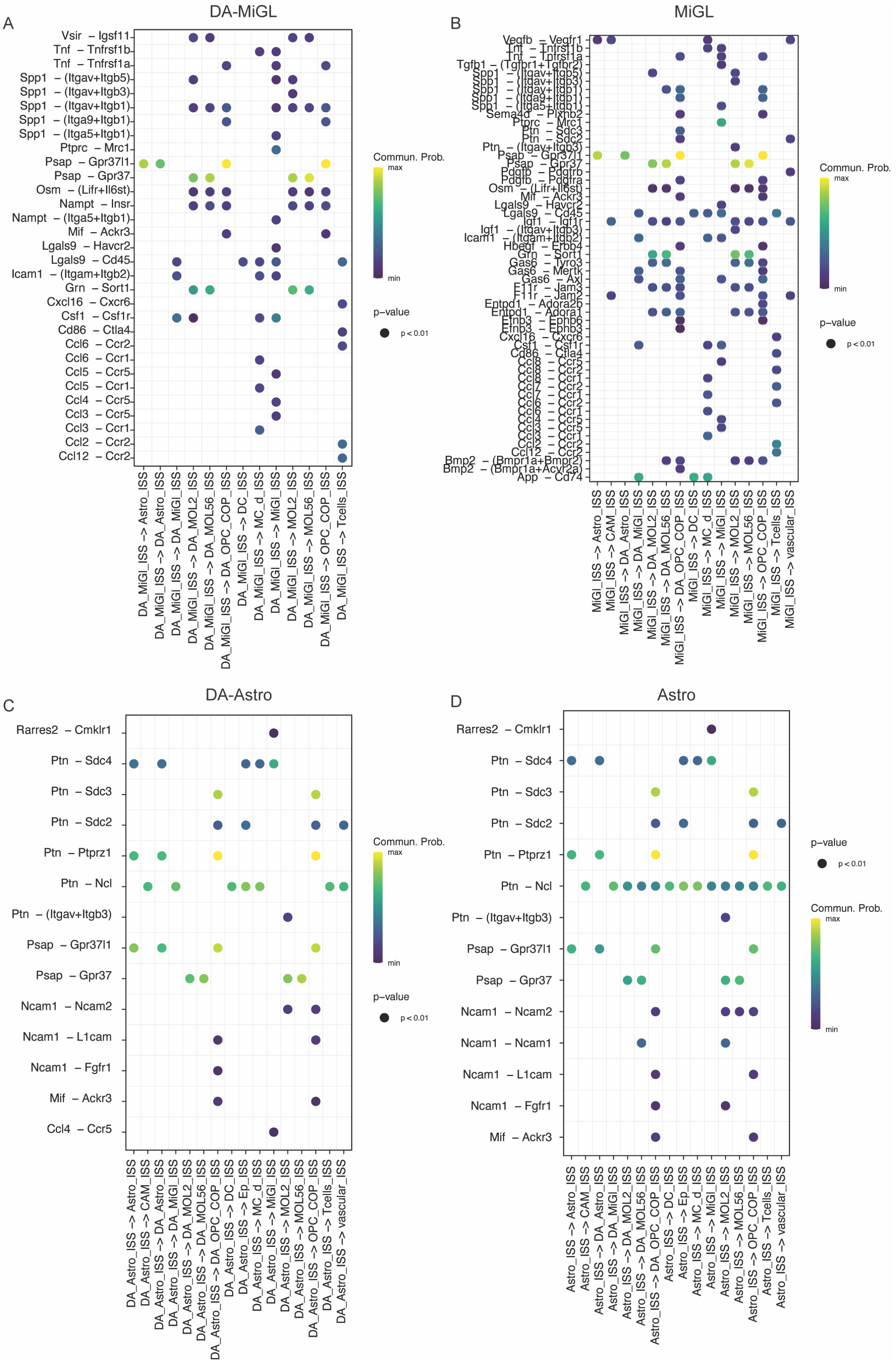

A
DA-MiGL
Vsir – Igsf11
Tnf – Tnfrsf1b
Tnf – Tnfrsf1a
Spp1 – (Itgav+Itgb5)
Spp1 – (Itgav+Itgb3)
Spp1 – (Itgav+Itgb1)
Spp1 – (Itga9+Itgb1)
Spp1 – (Itga5+Itgb1)
Ptprc – Mrc1
Psap – Gpr37l1
Psap – Gpr37
Osm – (Lifr+Il6st)
Nampt – Insr
Nampt – (Itga5+Itgb1)
Mif – Ackr3
Lgals9 – Havcr2
Lgals9 – Cd45
Icam1 – (Itgam+Itgb2)
Grn – Sort1
Cxcl16 – Cxcr6
Csf1 – Csf1r
Cd86 – Ctla4
Ccl6 – Ccr2
Ccl6 – Ccr1
Ccl5 – Ccr5
Ccl5 – Ccr1
Ccl4 – Ccr5
Ccl3 – Ccr5
Ccl3 – Ccr1
Ccl2 – Ccr2
Ccl12 – Ccr2
DA_MiGl_ISS –> Astro_ISS
DA_MiGl_ISS –> DA_Astro_ISS
DA_MiGl_ISS –> DA_MiGl_ISS
DA_MiGl_ISS –> DA_MOL2_ISS
DA_MiGl_ISS –> DA_MOL56_ISS
DA_MiGl_ISS –> DA_OPC_COP_ISS
DA_MiGl_ISS –> DC_ISS
DA_MiGl_ISS –> MC_d_ISS
DA_MiGl_ISS –> MiGl_ISS
DA_MiGl_ISS –> MOL2_ISS
DA_MiGl_ISS –> MOL56_ISS
DA_MiGl_ISS –> OPC_COP_ISS
DA_MiGl_ISS –> Tcells_ISS
Commun. Prob.
max
min
p–value
p < 0.01
B
MiGL
Vegfb – Vegfr1
Tnf – Tnfrsf1b
Tnf – Tnfrsf1a
Tgfb1 – (Tgfbr1+Tgfbr2)
Spp1 – (Itgav+Itgb5)
Spp1 – (Itgav+Itgb3)
Spp1 – (Itgav+Itgb1)
Spp1 – (Itga9+Itgb1)
Spp1 – (Itga5+Itgb1)
Sema4d – Plxnb2
Ptprc – Mrc1
Ptn – Sdc3
Ptn – Sdc2
Ptn – (Itgav+Itgb3)
Psap – Gpr37l1
Psap – Gpr37
Pdgfb – Pdgfrb
Pdgfb – Pdgfra
Osm – (Lifr+Il6st)
Mif – Ackr3
Lgals9 – Havcr2
Lgals9 – Cd45
Igf1 – Igf1r
Igf1 – (Itgav+Itgb3)
Icam1 – (Itgam+Itgb2)
Hbegf – Erbb4
Grn – Sort1
Gas6 – Tyro3
Gas6 – Mertk
Gas6 – Axl
F11r – Jam3
F11r – Jam2
Entpd1 – Adora2b
Entpd1 – Adora1
Efnb3 – Ephb6
Efnb3 – Ephb3
Cxcl16 – Cxcr6
Csf1 – Csf1r
Cd86 – Ctla4
Ccl8 – Ccr5
Ccl8 – Ccr2
Ccl8 – Ccr1
Ccl7 – Ccr2
Ccl7 – Ccr1
Ccl6 – Ccr1
Ccl4 – Ccr5
Ccl3 – Ccr5
Ccl3 – Ccr1
Ccl2 – Ccr2
Ccl12 – Ccr2
Bmp2 – (Bmpr1a+Bmpr2)
Bmp2 – (Bmpr1a+Acvr2a)
App – Cd74
MiGl_ISS –> Astro_ISS
MiGl_ISS –> CAM_ISS
MiGl_ISS –> DA_Astro_ISS
MiGl_ISS –> DA_MiGl_ISS
MiGl_ISS –> DA_MOL2_ISS
MiGl_ISS –> DA_MOL56_ISS
MiGl_ISS –> DA_OPC_COP_ISS
MiGl_ISS –> DC_ISS
MiGl_ISS –> MC_d_ISS
MiGl_ISS –> MiGl_ISS
MiGl_ISS –> MOL2_ISS
MiGl_ISS –> MOL56_ISS
MiGl_ISS –> OPC_COP_ISS
MiGl_ISS –> Tcells_ISS
MiGl_ISS –> vascular_ISS
Commun. Prob.
max
min
p–value
p < 0.01
C
DA-Astro
Rarres2 – Cmklr1
Ptn – Sdc4
Ptn – Sdc3
Ptn – Sdc2
Ptn – Ptprz1
Ptn – Ncl
Ptn – (Itgav+Itgb3)
Psap – Gpr37l1
Psap – Gpr37
Ncam1 – Ncam2
Ncam1 – L1cam
Ncam1 – Fgfr1
Mif – Ackr3
Ccl4 – Ccr5
DA_Astro_ISS –> Astro_ISS
DA_Astro_ISS –> CAM_ISS
DA_Astro_ISS –> DA_Astro_ISS
DA_Astro_ISS –> DA_MiGl_ISS
DA_Astro_ISS –> DA_MOL2_ISS
DA_Astro_ISS –> DA_MOL56_ISS
DA_Astro_ISS –> DA_OPC_COP_ISS
DA_Astro_ISS –> DC_ISS
DA_Astro_ISS –> Ep_ISS
DA_Astro_ISS –> MC_d_ISS
DA_Astro_ISS –> MiGl_ISS
DA_Astro_ISS –> MOL2_ISS
DA_Astro_ISS –> MOL56_ISS
DA_Astro_ISS –> OPC_COP_ISS
DA_Astro_ISS –> Tcells_ISS
DA_Astro_ISS –> vascular_ISS
Commun. Prob.
max
min
p–value
p < 0.01
D
Astro
Rarres2 – Cmklr1
Ptn – Sdc4
Ptn – Sdc3
Ptn – Sdc2
Ptn – Ptprz1
Ptn – Ncl
Ptn – (Itgav+Itgb3)
Psap – Gpr37l1
Psap – Gpr37
Ncam1 – Ncam2
Ncam1 – Ncam1
Ncam1 – L1cam
Ncam1 – Fgfr1
Mif – Ackr3
Astro_ISS –> Astro_ISS
Astro_ISS –> CAM_ISS
Astro_ISS –> DA_Astro_ISS
Astro_ISS –> DA_MiGl_ISS
Astro_ISS –> DA_MOL2_ISS
Astro_ISS –> DA_MOL56_ISS
Astro_ISS –> DA_OPC_COP_ISS
Astro_ISS –> DC_ISS
Astro_ISS –> Ep_ISS
Astro_ISS –> MC_d_ISS
Astro_ISS –> MiGl_ISS
Astro_ISS –> MOL2_ISS
Astro_ISS –> MOL56_ISS
Astro_ISS –> OPC_COP_ISS
Astro_ISS –> Tcells_ISS
Astro_ISS –> vascular_ISS
p–value
p < 0.01
Commun. Prob.
max
min

***Supplementary Figure 8, Receptor ligand interactions for DA-microglia and DA-astrocytes in EAE, related to Figure 5.***

***(A)*** *Receptor ligand interactions for DA-MiGL with DA-MiGL as a source.*

***(B)*** *Receptor ligand interactions for MiGL with MiGL as a source.*

***(C)*** *Receptor ligand interactions for DA-Astro with DA-Astro as a source.*

***(D)*** *Receptor ligand interactions for Astro with Astro as a source.*

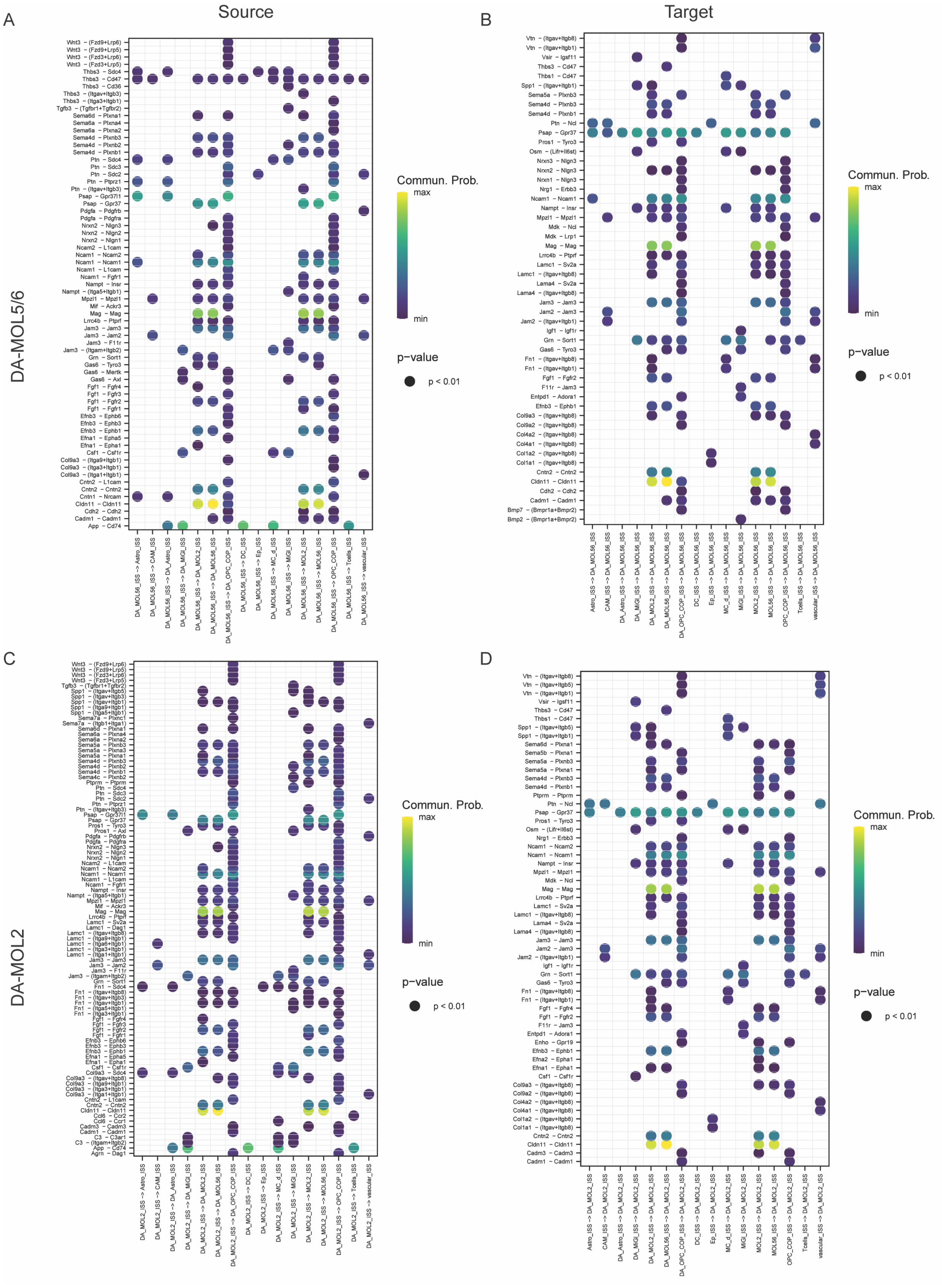

A
Source
B
Target
C
D
DA-MOL5/6
DA-MOL2
Commun. Prob.
max
min
p-value
p < 0.01

***Supplementary Figure 9, Receptor ligand interactions for DA MOLs in EAE, related to Figure 5.***

***(A)*** *Receptor ligand interactions for DA-MOL5/6 with DA-MOL5/6 as a source.*

***(B)*** *Receptor ligand interactions for DA-MOL5/6 with DA-MOL5/6 as a target.*

***(C)*** *Receptor ligand interactions for DA-MOL2 with DA-MOL2 as a source.*

***(D)*** *Receptor ligand interactions for DA-MOL2 with DA-MOL2 as a target.*

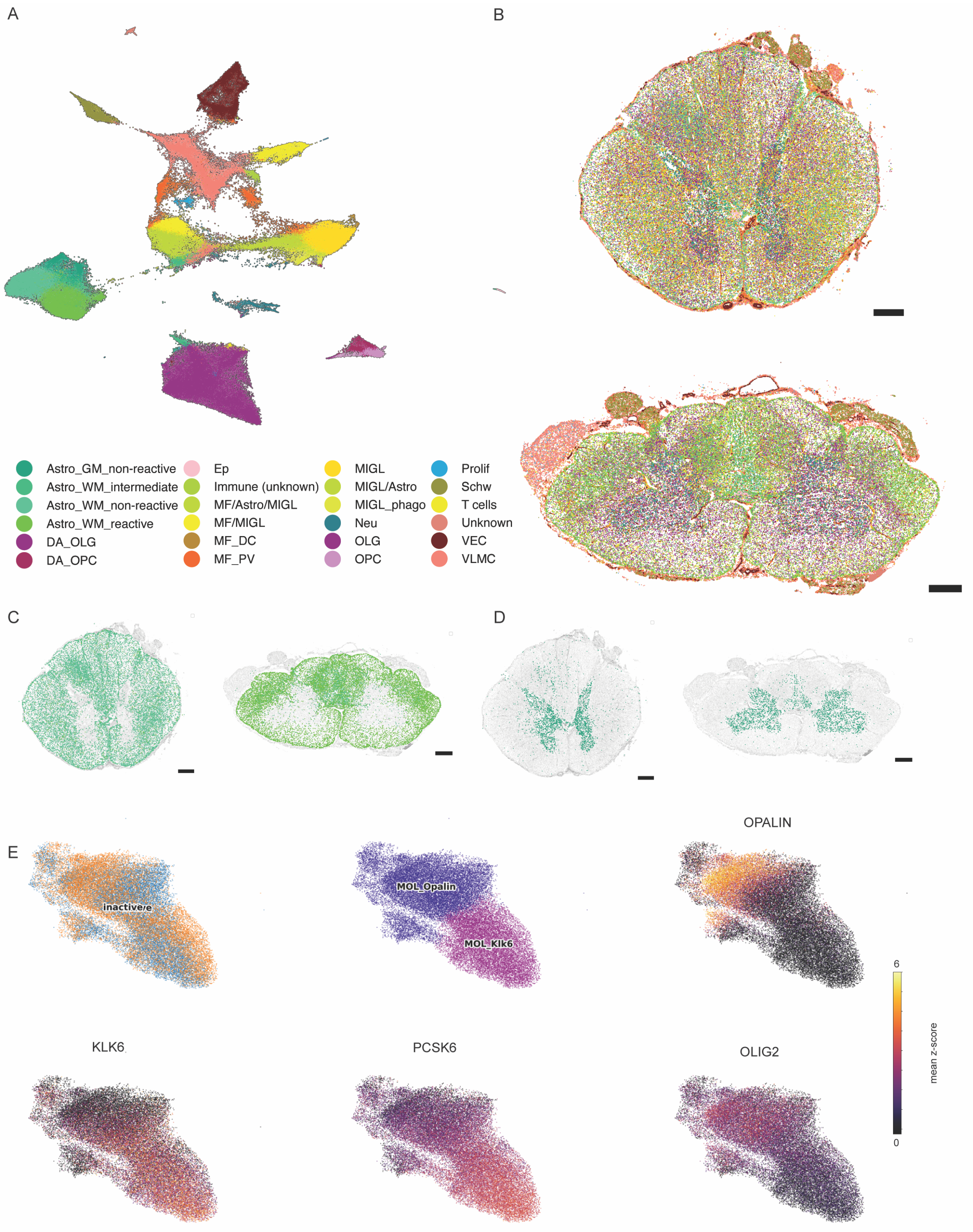
A
B
Astro_GM_non-reactive
Astro_WM_intermediate
Astro_WM_non-reactive
Astro_WM_reactive
DA_OLG
DA_OPC
Ep
Immune (unknown)
MF/Astro/MIGL
MF/MIGL
MF_DC
MF_PV
MIGL
MIGL/Astro
MIGL_phago
Neu
OLG
OPC
Prolif
Schw
T cells
Unknown
VEC
VLMC
C
D
E
inactivere
MOL_Opalin
MOL_Klk6
OPALIN
KLK6
PCSK6
OLIG2
6
0
mean z-score

***Supplementary Figure 10, Cell populations identified by ISS in human MS cervical spinal cord, related to Figure 6.***

***(A)*** *UMAP of cell types annotated using hierarchical clustering, intermediate level of cell type annotation.*

***(B)*** *Spatial maps of cell types. The colours of the cell types correspond to the colours in* ***A.*** *Scale bars - 1000µm.*

***(C)*** *White matter astrocytes, colours correspond to* ***A.*** *Scale bars - 1000µm.*

***(D)*** *Grey matter astrocytes, colours correspond to* ***A.*** *Scale bars - 1000µm.*

***(E)*** *UMAPs of mature oligodendrocytes coloured by the sample type, annotation, and expression of OPALIN, KLK6, PCSK6 and OLIG2.*

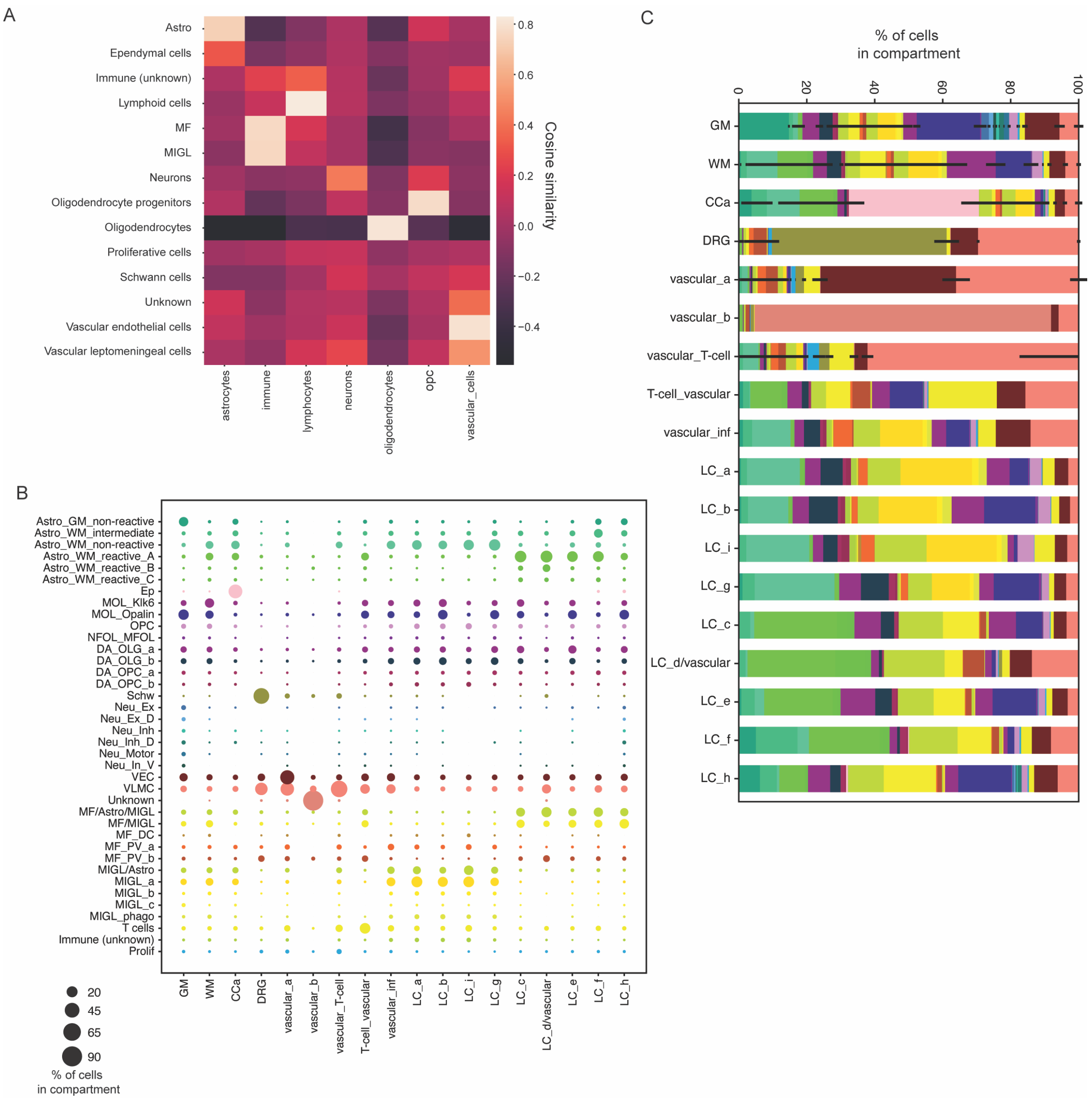


***Supplementary Figure 11, Composition of compartments identified by ISS in human MS cervical spinal cord, related to Figure 7.***

***(A)*** *Cosine similarity between broad annotated cell types in our human MS data and cell types from Absinta et al. 2021.*

***(B)*** *Dot plot representing the compartment preference of cell types.*

***(C)*** *Stacked histogram showing the relative compartment composition, normalised by the total number of cells in that compartment.*
