## Supplementary Tables for "Single cell-resolution *in situ* sequencing elucidates spatial dynamics of multiple sclerosis lesion and disease evolution"

**Supplementary Table 1**. Genes used in the mouse experiments.

| Gene |
|---|
| 2610035D17Rik |
| 9630013A20Rik |
| Adora2b |
| Aif1 |
| Aldh1l1 |
| Anln |
| Anxa5 |
| Anxa8 |
| Apod |
| Aqp4 |
| Arc |
| Arrdc2 |
| B2m |
| Batf3 |
| Birc2 |
| Bmp4 |
| Brca1 |
| Btg2 |
| C030029H02Rik |
| C4b |
| Cacna1d |
| Car2 |
| Cavin1 |
| Cbr2 |
| Ccl17 |
| Ccl2 |
| Ccl5 |
| Ccr2 |
| Ccr7 |
| Cd163 |
| Cd19 |
| Cd209a |
| Cd24a |
| Cd3g |
| Cd4 |
| Cd44 |

| |
|---|
| Cd59a |
| Cd69 |
| Cd74 |
| Cd8a |
| Cdkn1a |
| Cdkn1c |
| Cemip2 |
| Chat |
| Ciita |
| Cldn10 |
| Cldn11 |
| Clec9a |
| Clic4 |
| Cnksr3 |
| Col20a1 |
| Crip1 |
| Cspg5 |
| Ctla4 |
| Ctps |
| Ctss |
| Cx3cr1 |
| Ddr1 |
| Dnajb1 |
| Dusp1 |
| Ednrb |
| Egr1 |
| Egr2 |
| Fam107a |
| Fam107b |
| Fam214a |
| Fcgr2b |
| Fcrls |
| Flt3 |
| Fos |
| Fosb |
| Foxp3 |
| Frmd4a |
| Fyn |

| |
|---|
| Gad1 |
| Gad2 |
| Gbp7 |
| Gfap |
| Gja1 |
| Gjb1 |
| Glul |
| Gng11 |
| Gpr34 |
| Gpr37l1 |
| Grm3 |
| H2-Aa |
| H2-Ab1 |
| H2-D1 |
| H2-DMa |
| H2-Eb1 |
| H2-K1 |
| H2-T23 |
| Hapln2 |
| Hcn2 |
| Hexb |
| Hif3a |
| Hopx |
| Hspa1a |
| Icos |
| Ifih1 |
| Ifit1 |
| Ighm |
| Igtp |
| Il12rb1 |
| Il33 |
| Irf7 |
| Irf8 |
| Irgm1 |
| Irgm2 |
| Itga8 |
| Itgae |
| Itgam |

| |
|---|
| Itgax |
| Itih5 |
| Itpr2 |
| Jcad |
| Kcnip3 |
| Klf4 |
| Klk6 |
| Klk8 |
| Lat |
| Lbh |
| Lgals1 |
| Lrp1 |
| Ly6c1/2 |
| Ly86 |
| Lyve1 |
| Mal |
| Mbp |
| Mcam |
| Meg3 |
| Mertk |
| Mgst3 |
| Mki67 |
| Mog |
| Mpzl1 |
| Mrc1 |
| Ms4a1 |
| Ms4a7 |
| Mt1 |
| Mt2 |
| Mycl |
| Mylk |
| Ninj2 |
| Nlrc5 |
| Nr4a1 |
| Nrn1 |
| Olfml3 |
| Olig2 |
| Onecut2 |

| |
|---|
| Opalin |
| P2ry12 |
| P2ry13 |
| Panx1 |
| Pcdh15 |
| Pdcd1 |
| Pdgfra |
| Pdgfrb |
| Pf4 |
| Phyhd1 |
| Plac8 |
| Plin4 |
| Pllp |
| Pmp22 |
| Prr5l |
| Psmb8 |
| Psmb9 |
| Ptgds |
| Ptprz1 |
| Qdpr |
| Rab37 |
| Rap2a |
| Rgs20 |
| Rnase4 |
| Rorgt |
| Rph3a |
| Rras2 |
| S100a10 |
| S100b |
| Sdc1 |
| Selenop |
| Selplg |
| Sema4d |
| Serpina3c |
| Serpina3f |
| Serpina3h |
| Serpina3i |
| Serpina3m |

| |
|---|
| Serpina3n |
| Serpind1 |
| Serpine2 |
| Serping1 |
| Sgk1 |
| Sh3bp4 |
| Siglec1 |
| Siglech |
| Slc14a1 |
| Slc17a6 |
| Slc17a7 |
| Slc1a1 |
| Slc32a1 |
| Slc5a7 |
| Slc6a1 |
| Slc6a5 |
| Slc9a3r2 |
| Slco3a1 |
| Slk |
| Sox10 |
| Sox6 |
| Sox8 |
| Spock3 |
| Stab1 |
| Stat1 |
| Stmn2 |
| Sult1a1 |
| Syt4 |
| Tap1 |
| Tbx21 |
| Thbs3 |
| Tlr3 |
| Tmem119 |
| Tmem163 |
| Tmem176a |
| Tnfrsf1a |
| Tnr |
| Trbc1 |

| |
|---|
| Trem2 |
| Trim34a |
| Tssc4 |
| Ttr |
| Ttyh1 |
| Tubb3 |
| Vcan |
| Vtn |
| Vxn |
| Wfdc18 |
| Zbp1 |
| Zbtb46 |
| Zfand4 |

**Supplementary Table 2**. Genes used in the human experiments.

| Gene |
|---|
| ABCC9 |
| ADAMTS12 |
| ADAMTS16 |
| ADAMTS3 |
| ADRA1A |
| ADRA1B |
| AIF1 |
| ALK |
| ANGPT1 |
| ANK1 |
| ANKRD18A |
| ANO3 |
| ANXA1 |
| APOE |
| APP |
| AQP4 |
| ARHGAP24 |
| ATP2C2 |
| B4GALNT1 |
| BCAN |
| BRINP3 |
| BTBD11 |
| C1QL3 |
| C1orf162 |
| CALCRL |
| CAPG |
| CAPN3 |
| CAV1 |
| CCK |
| CCL4 |
| CCL5 |
| CCNA1 |
| CCNB2 |
| CD14 |
| CD163 |
| CD2 |
| CD36 |
| CD3G |
| CD4 |
| CD48 |
| CD52 |
| CD68 |
| CD83 |
| CD86 |
| CDH1 |
| CDH12 |

| |
|---|
| CDH4 |
| CDH6 |
| CDK1 |
| CEMIP |
| CEMIP2 |
| CENPF |
| CHODL |
| CLDN11 |
| CNDP1 |
| CNTN2 |
| CNTNAP3B |
| COL12A1 |
| COL25A1 |
| CORO1A |
| CRHBP |
| CRYM |
| CSPG4 |
| CTNNA3 |
| CTSH |
| CTSS |
| CUX2 |
| CX3CR1 |
| CXCL14 |
| CXCR4 |
| CYTIP |
| DCN |
| DDR2 |
| DNER |
| EFHD1 |
| EGFR |
| ELOVL2 |
| ERBB3 |
| ERMN |
| EYA4 |
| FASLG |
| FBLN1 |
| FCER1G |
| FCGR1A |
| FCGR3A |
| FGFR2 |
| FGFR3 |
| FILIP1 |
| FLT1 |
| FSTL4 |
| GAD1 |
| GAD2 |
| GAS2L3 |
| GJA1 |
| GNLY |

| |
|---|
| GPNMB |
| GPR183 |
| GPR34 |
| GZMA |
| HES1 |
| HHATL |
| HILPDA |
| HLA-DMB |
| HLA-DQA1 |
| HS3ST2 |
| HS3ST4 |
| HTR2A |
| HTR2C |
| IDH1 |
| IDH2 |
| IDO1 |
| IFITM3 |
| IGFBP3 |
| IGFBP4 |
| IGFBP5 |
| IL7R |
| ITGA8 |
| ITGAM |
| ITGAX |
| ITGB2 |
| KCNH5 |
| KIT |
| KLF2 |
| KLF4 |
| KLK6 |
| KLRB1 |
| LAMA2 |
| LAMP5 |
| LHX6 |
| LOX |
| LRRK1 |
| LRRK2 |
| LY86 |
| LYPD6 |
| LYPD6B |
| LYVE1 |
| MAG |
| MAL |
| MCTP2 |
| MEIS2 |
| MEPE |
| MGST1 |
| MKI67 |
| MOBP |
| MOG |

| |
|---|
| MS4A6A |
| MYO16 |
| MYO5B |
| MYRF |
| NCSTN |
| NDST4 |
| NES |
| NKG7 |
| NNAT |
| NOTCH1 |
| NPFFR2 |
| NPNT |
| NPY1R |
| NR2F2 |
| NR4A2 |
| NRP1 |
| NTNG1 |
| NTNG2 |
| NWD2 |
| NXPH2 |
| OLIG1 |
| OLIG2 |
| OPALIN |
| OTOGL |
| P2RY12 |
| P2RY13 |
| PAX6 |
| PCNA |
| PCSK1 |
| PCSK6 |
| PDGFD |
| PDGFRA |
| PECAM1 |
| PHLDB2 |
| PLCE1 |
| PLCH1 |
| PLD5 |
| POSTN |
| POU6F2 |
| PROX1 |
| PSEN1 |
| PSEN2 |
| PSENEN |
| PTCHD4 |
| PTPRC |
| PTPRZ1 |
| PVALB |
| RASGRP1 |
| RELN |
| RFTN1 |

| |
|---|
| RGS10 |
| RGS16 |
| RIT2 |
| RNASET2 |
| RNF144B |
| RORB |
| ROS1 |
| RSPO2 |
| RXFP1 |
| RYR3 |
| S100A4 |
| SAMD5 |
| SDK1 |
| SEMA5A |
| SERPINA3 |
| SFRP2 |
| SLC17A6 |
| SLC17A7 |
| SLC24A3 |
| SLC26A4 |
| SLIT3 |
| SNCG |
| SNTB2 |
| SORCS1 |
| SOX10 |
| SOX11 |
| SOX2 |
| SOX4 |
| SOX9 |
| SPHKAP |
| SPI1 |
| SPON1 |
| SST |
| ST18 |
| STAT3 |
| STK32B |
| STXBP2 |
| SULF1 |
| SYNPR |
| TAC1 |
| TACR1 |
| TENM1 |
| TESPA1 |
| TGFB1 |
| TGFB2 |
| TGFBI |
| THBS1 |
| THEMIS |
| THSD4 |
| THSD7B |

| |
|---|
| TMEM132C |
| TMIGD3 |
| TOP2A |
| TP53 |
| TPH2 |
| TRAC |
| TREM2 |
| TRHDE |
| TRIL |
| TRPC5 |
| TRPC6 |
| TSHZ2 |
| TTYH1 |
| UGT8 |
| UNC5B |
| VCAN |
| VIP |
| VWC2L |
| WIF1 |
| ZBBX |
| ZDHHC23 |
